# Extracellular vesicles mediate the intercellular exchange of nanoparticles

**DOI:** 10.1101/2021.02.23.432325

**Authors:** Xian Wu, Tang Tang, Yushuang Wei, Katherine A. Cummins, David K. Wood, Hong-Bo Pang

## Abstract

In order to exert their therapeutic effects, nanoparticles (NPs) often need to travel into the tissues composed of multilayered cells. Accumulative evidence has revealed the crucial role of transcellular transport route (entry into one cell, exocytosis, and re-entry into another) in this process. While NP endocytosis and subcellular transport have been intensively characterized, the exocytosis and re-entry steps are poorly understood, which becomes a barrier to improve NP delivery into complex tissues. Here, we termed the exocytosis and re-entry steps together as intercellular exchange. We developed a collagen-based 3D assay to specifically monitor and quantify the intercellular exchange events of NPs, and distinguish the contributions of several potential mechanisms. Our results showed that NPs can be exocytosed freely or enclosed inside extracellular vesicles (EVs) for re-entry, while direct cell-cell contact is hardly involved. EVs account for a significant fraction of NP intercellular exchange, and its importance in NP transport was demonstrated *in vitro* and *in vivo*. Intriguingly, while freely released NPs engage with the same cellular receptors for re-entry, EV-enclosed ones bypass this dependence. These studies provide an easy and precise system to investigate the intercellular exchange stage of NP delivery, and shed the first light in the importance of EVs in NP transport between cells and across complex tissues.

## Introduction

Due to the tunable physicochemical characteristics and versatile cargo loading properties, nanoparticles (NPs) have great potential to improve the diagnosis and treatment of human diseases^1–5^. One prerequisite for many *in vivo* applications of NPs is to travel efficiently in the tissue of multilayered cells, and eventually into target cells. Therefore, understanding the cell biology, especially the transport pathways, is of particular significance to the success of nanomedicine. Over the years, numerous efforts have been devoted to elucidate how NPs of various types enter the cell and travel inside^6–8^. However, fewer studies have focused on another fundamental question: how will these internalized NPs be released from one cell, and transferred to another?

Besides being a fundamental cellular process, this question is also of great relevance to NP translation into the clinics. Since the first NP-formulated drug (Doxil) was approved in 1995^9^, the rate of NP clinical translation has been limited. A major challenge has been the poor delivery efficiency into solid organs or tissues composed of multilayered cells^10–12^. This problem is best exemplified in solid tumors in which studies have shown that only ~0.7% (median) of systemically injected dosage of NPs eventually accumulate in the tumor tissue^13^. Traditionally, the central paradigm of NP transport into tumors was the enhanced permeability and retention (EPR) effect, which considered passive diffusion through intercellular gaps as the primary route for extravasation. Recently, a series of studies showed that the majority of NPs rather extravasate through an active transcellular transport pathway. Using transmission electron microscopy (TEM), Sindhwani et al. found that tumor endothelium is largely intact without gaps, and NPs mainly reside inside endothelial cells during extravasation^14^. Using a fixation-based method, the authors also inactivated the active transport process prior to NP administration and observed that this treatment eliminates the majority of NP extravasation and tumor accumulation. Another line of evidence arises from the studies on a tumor-penetrating peptide, iRGD. This peptide can actively penetrate across tumor vessels and deeply into the extravascular regions when covalently coupled to various cargo types, ranging from small molecules to NPs^15^. This process is also energy dependent, and the penetration distance is far beyond the capability of passive diffusion^15^. It was later shown that iRGD-coupled NPs also reside inside endothelial cells during extravasation^16^. Together, these results indicate that NPs need to first enter endothelial cells, and then be exported for entering subsequent cells. This highlights the central role of the active (energy dependent) transcellular route in NP delivery and *in vivo* applications.

Theoretically, transcellular transport consists of four steps: entry into one cell, intracellular transport, cargo export or exocytosis, and the re-entry of released cargo into a second cell. While the first two steps of NP transport have been well characterized, the latter two remain largely understudied^17,18^. Here, we term the last two stages (cargo export and re-entry) as intercellular exchange. Traditional studies on NP transcytosis cover exocytosis but not re-entry^7^, and so far, there is no assay specifically quantifying this process. Therefore, there was a need for assays that monitor and quantify the intercellular exchange events of NPs independent of interference from the first two steps of transcellular transport. We previously developed one such assay, integrating cell-penetrating peptides (CPPs) with a novel class of etchable NPs. Our results showed that a significant number of NPs are transferred from one cell (donor) to another (recipient) in membrane-enclosed structures, which we hypothesized to be either secreted extracellular vesicles (EVs) or through direct cell-cell contact^19^. Here, we adapted this assay into a collagen-based format in three dimensions (3D) to further elucidate the underlying mechanism. The current format is easy to set up, better mimics the cell growth environment *in vivo*, and is capable of distinguishing several possible routes for intercellular exchange. Our results here demonstrated that EVs, but not direct cell-cell contact, serve as the membrane-enclosed conduit for intercellular exchange of NPs. The EV route accounted for a significant and varying fraction of intercellular exchange, and its importance was proven *in vitro* and *in vivo*. Our study also unveiled differences for freely released and EV-carrying NPs to enter the recipient cells.

## Results

### A 3D intercellular exchange assay

The intercellular exchange assay was established as below. First, donor cells were incubated with CPP-functionalized NPs for internalization. The primary donor cell types included human umbilical vein endothelial cells (HUVECs) and PC-3 (human prostate cancer cell line), as we aim to understand the cargo transfer from endothelium to other cell types, as well as the material exchange between cells beyond the vasculature. The key to our assay is etchable silver-based NPs (AgNPs). In etching, a chemical and nontoxic solution, etchant, is used to rapidly dissolve AgNPs and thus eliminate their fluorescence signals^19,20^. Importantly, etchant cannot permeate lipid membranes, such as cell membranes. Therefore, etching can remove extracellular and cell surface bound AgNPs, but those internalized remain intact. By eliminating these “noise” signals caused by extracellular NPs, etching is particularly useful in improving the quality of cellular and *in vivo* imaging ^20^. To facilitate their entry into donor cells, we functionalized AgNPs with two CPPs. Transactivating transcriptional activator (TAT) is the first and one of the most widely used CPPs to deliver NPs and macromolecules into cells^21–23^. RPARPAR is the prototype of a novel class of CPPs, CendR peptides, whose cellular receptor is neuropilin-1 (NRP1) ^24^. iRGD is a tumor-specific CendR peptide, and NRP1 binding is the basis for its vascular and tumor penetration property ^15,25^. We previously showed that these peptides, while engaging with different cellular receptors, invoke a similar macropinocytosis-like process for cell entry^19,26^. Besides assisting the NP uptake, these CPPs were also used to lead NPs from one cell to another^16,27^, and may help elucidate the function of ligand-receptor interactions during intercellular exchange.

After CPP-AgNP internalization, donor cells were etched to ensure that AgNPs reside only inside the cells. Then, these cells were lifted up and mixed with another group of cells, termed recipient cells. Recipient cells are labeled in a different fluorescence color, so that we can distinguish these two cell types. After incubation, the cell mixture was dissociated into single cells, and AgNPs in donor and recipient cells were detected and quantified by flow cytometry (Fig. S1). Due to etching, AgNPs in recipient cells can only arise from the exocytosis from donor and then re-entry into recipient cells, thus the intercellular exchange. In all our studies, donor cells were incubated with CPP-AgNPs and etched in bulk, and then evenly aliquoted into different experimental groups. This way, we ensure that the entry and subcellular transport status of AgNPs in donor cells are all the same, thus decoupling these two steps from the quantification of intercellular exchange. Plus, this allows us to simplify the analysis, and directly define the intercellular exchange efficiency as the total AgNPs that can reach recipient cells.

The first version of our assay simply cultured the donor/recipient cell mixture in ultra-low binding plates, which is vastly different from the physiological conditions of cell growth^19^. Here, we adapted it to a 3D format. We encapsulated recipient cells in a collagen matrix on the bottom of the wells in the plate, and seeded CPP-NP-containing donor cells on top of the recipient-collagen layer (Fig. 1A). PC-3 and PC3-GFP cells were first used as donor and recipient cells, respectively, and TAT-coated AgNPs (T-AgNPs) and RPARPAR-coated AgNPs (R-AgNPs) as the model CPP-NPs for assay optimization. We confirmed that constant etching in this system induced little change on cell viability (Fig. S2). We observed ~9 % of recipient cells becoming positive for both T-AgNPs and R-AgNPs (Fig. 1B), and the intercellular exchange efficiency was defined accordingly (see Methods). Several parameters were further optimized to maximize the intercellular exchange, such as cell number, donor/recipient ratio, the effect of fetal bovine serum in the medium and incubation time (Fig. S3). The detailed process of optimization for the experimental condition is described in supplementary information (Fig. S3), and used throughout this study. Interestingly, we observed little difference in the intercellular exchange efficiency when plating donor cells on top either as monolayer or when encapsulated in a separate collagen layer (Fig. S3E). For simplicity, we used the monolayer format for donor cells in the following experiments.

**Fig. 1.**
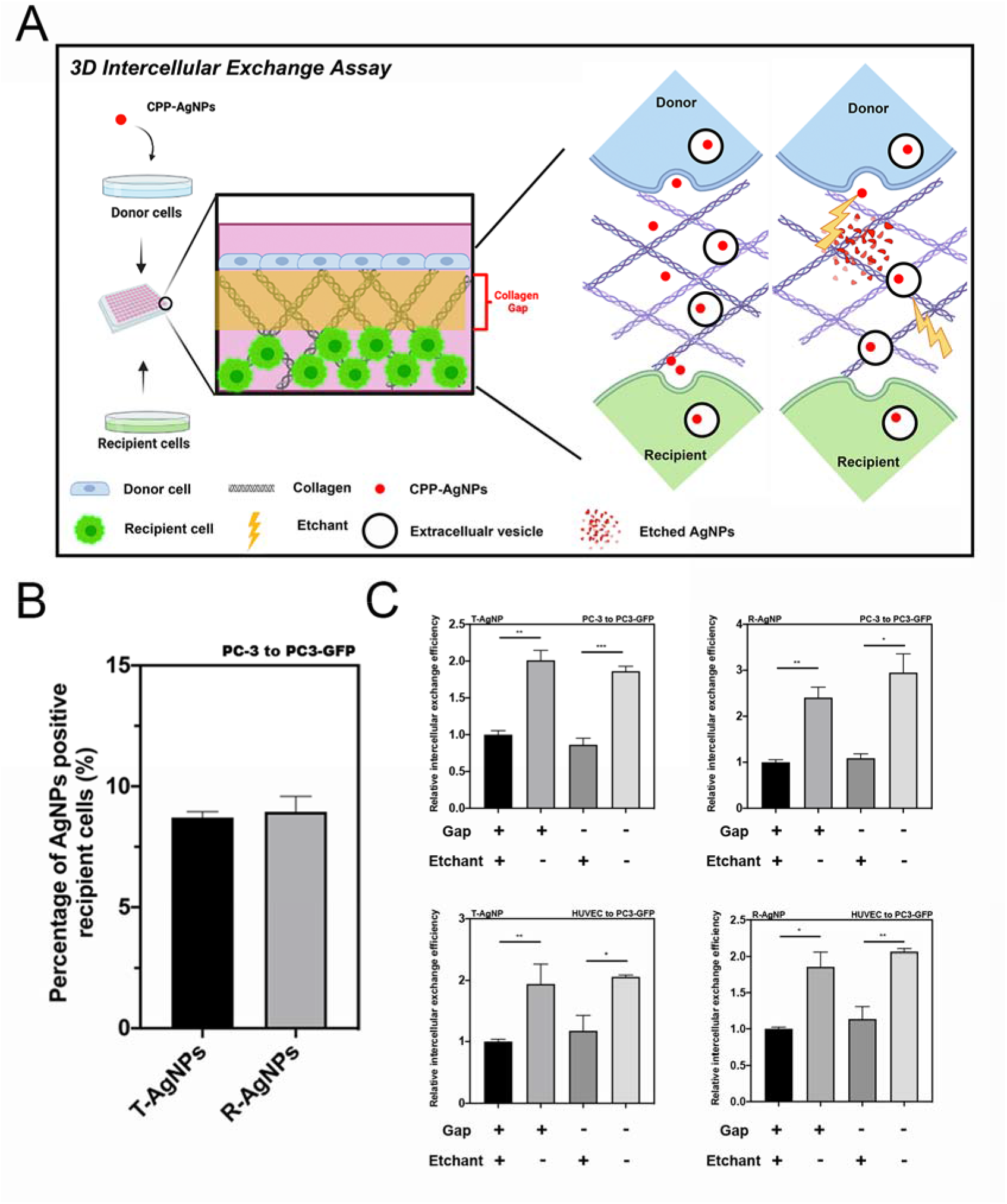
Intercellular exchange of CPP-AgNPs were monitored and quantified by 3D intercellular exchange assay. (A) Schematic illustration of the 3D intercellular exchange assay used throughout this study and EV-mediated penetration. (B) Percentage of CPP-AgNPs positive recipient cells in the intercellular exchange assay. Intercellular exchange assay of T-AgNPs and R-AgNPs from PC-3 to PC3-GFP cells was carried out as described in Methods, respectively. The percentage of CPP-AgNPs positive recipient cells was quantified by flow cytometry analysis (y axis). (C) The 3D intercellular exchange assays with different pairs of donor/recipient cells. After 24 h incubation under the indicated conditions (x axis), intercellular exchange efficacy of T-AgNPs / R-AgNPs from PC-3 to PC3-GFP / HUVEC to PC3-GFP cells was quantified as described in Methods and normalized to that of under constant etching with collagen gap (y axis), respectively. Error bars indicate S.E.M., n=3. *P<0.05, **P<0.01 and ***P<0.001 (Student’s t-test).

### Extracellular vesicles, but not direct cell-cell contact, mediate the NP exchange between cells

Besides better mimicking the native growth environment, the 3D assay is of unique advantage to distinguish several possible routes for intercellular exchange. We speculate three possible routes of intercellular exchange: CPP-NPs are exported as free agents (direct release), or are exocytosed into EVs, or are exchanged via direct contact between donor and recipient cells. For the first route, CPP-NPs are directly exposed to surrounding environment, and will recognize and bind to the same receptor on recipient cells as donor cells. Alternatively, CPP-NPs will be protected from etching by lipid membranes in the latter two routes. Thus, constant etching of donor/recipient mixtures can distinguish the direct release from the other two routes, while it failed to distinguish between EV release and direct cell-cell contact. To solve this problem, an extra collagen gap (no cells) was added between donor and recipient cells to prevent cell-cell contact (Fig. 1A). We performed immune-histochemical and immunofluorescence staining on the 3D collagen matrix (Fig. S4), and confirmed that no cells or cellular structures were seen in the gap region 24 h after seeding cells in our system, thus no direct cell-cell contact. Using etchant and this gap together, we were able to distinguish the contributions from all three routes for intercellular exchange.

We first quantified the intercellular exchange efficiency of T-AgNPs and R-AgNPs in multiple donor/recipient pairs (Fig. 1C, Fig. S5 and S6). Constant etching caused a significant reduction of both CPP-AgNPs that can travel from donor to recipient cells, while the reduction range depends on CPP and donor/recipient cell types. This result suggests that a significant part of CPP-NPs is exported freely by donor cells. The collagen gap between donor and recipient cells as well as the concentration of collagen in the gap (Fig. S3C), however, exhibited little effect on the intercellular exchange of NPs, regardless of constant etching. We made the same observation regardless of the type of CPPs and donor/recipient cells used. This result indicates that direct cell-cell contact is not required for the etching-resistant intercellular exchange. Therefore, EVs are the etching-resistant carriers of CPP-NPs from donor to recipient cells. We further confirmed that etching treatment has little or no effect on total EV production, or EV uptake by donor/recipient cells (Fig. S7A-D) while efficient for eliminating free AgNPs (Fig. S7E).

The endocytic and exocytic efficiency of CPP-NPs were also measured. While we always supplied overwhelming NP concentration to donor cells for uptake, ~15-20% of total CPP-NPs input was taken up by donor cells after 4-h incubation, of which around 40% disappeared inside cells. It was reported that AgNPs remain intact and fluorescent up to 24-h inside cells^20^. Thus, this reduction was likely due to NP exocytosis in either free or EV-enclosed form (Fig. S8). AgNPs of different sizes were also used, which revealed a size dependence of intercellular exchange efficiency for T-AgNPs but not for R-AgNPs (Fig. S9).

### CPP-NPs are exocytosed inside EVs

EVs are a heterogenous group of secreted vesicles including exosomes and microvesicles ^28–30^. EVs have long been recognized as the conduit for intercellular communication and material exchange (e.g. nucleic acids, proteins)^31,32^. Here, we performed additional experiments to verify the notion that EVs mediate the intercellular exchange of NPs. First, we isolated EVs secreted by donor cells after CPP-NP internalization and etching. Using density gradient ultracentrifugation as previously described^33^, we were able to separate NP-containing from NP-free EVs (Fig. 2A). The density of NP-free EVs was find to be within the range of 1.127-1.136 g/mL, and this number for NP-containing EVs was 1.175-1.194 g/mL. Other than the visible separation of these two fractions, we also detected the fluorescence intensity of enclosed AgNPs to determine the purity of separation. Compared to NP-containing ones, NP-free EVs showed very little, if any, signals of AgNPs, supporting the notion that we were able to separate these two EV subsets (Fig. 2B). Next, we set out to investigate whether these AgNPs are inside the EVs. Free AgNPs released by cells were collected from the bottom fraction and the fluorescence of these AgNPs decreased significantly after etching due to the loss of antenna effect^20^ (Fig. 2C).

**Fig. 2.**
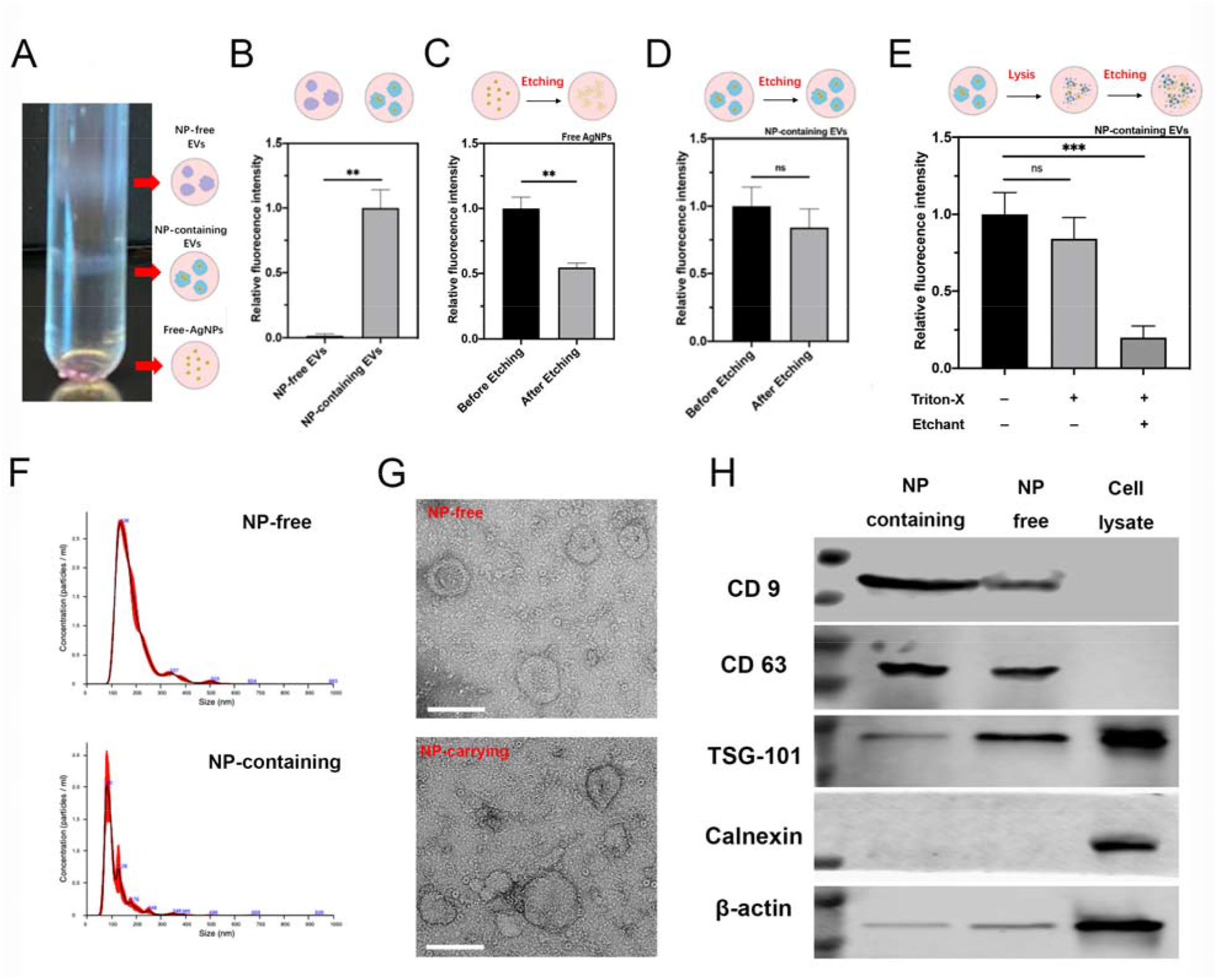
Validation of CPP-NPs exocytosed inside EVs. T-AgNPs were exocytosed by EVs. NP-free EVs and NP-containing EVs were isolated from PC-3 cells pre-incubated with T-AgNPs by density gradient ultracentrifugation and characterized as described in Methods. (A-E) T-AgNPs were released from cells as free NPs and EV-capsulated NPs. (A) Representative picture of fractions of NP-free EVs and NP-containing EVs after density gradient ultracentrifugation. (B) Fluorescence intensity of T-AgNPs in NP-free EVs and NP-containing EVs was detected and normalized to that of in NP-containing EVs (y axis). (C) Fraction containing free AgNPs was collected and the fluorescence intensity of T-AgNPs within before and after etching was detected and the result was normalized to that of before etching (y axis). (D) Fluorescence intensity of T-AgNPs within NP-containing EVs before and after etching was detected and the result was normalized to that of before etching (y axis). (E) After treating NP-containing EVs with TX-100 detergent, the fluorescence intensity of T-AgNPs within before / after etching was detected and the result was normalized to that of without any treatment (y axis). (F) Size distribution of NP-free and NP-containing EV detected by NTA (Red bars indicate S.D., n=3). (G) Representative TEM images of NP-free EVs and NP-containing EVs. Scale bar, 200nm. (H) Western blot analysis of NP-free and NP-containing EVs. The present of canonical exosome markers, including CD9, CD63 and TSG-101 and the absence of Calnexin were detected from NP-free EVs and NP-containing EVs. Error bars indicate S.E.M., n=3. **P<0.01, **** P<0.0001 and ns, no significance (Student’s t-test).

However, the fluorescence signal of AgNPs inside NP-containing EVs did not change significantly before and after etching (Fig. 2D). After lyzing the NP-containing EVs with detergent, the fluorescence signal of AgNPs significantly decreased after etching (Fig. 2E). All these data suggest that these exocytosed AgNPs are enclosed inside EVs.

We further analyzed the morphologic and molecular nature of these two EV subsets. Their mean diameter was around 155 nm for NP-containing EVs, and around 181 nm for NP-free ones measured by Nano Tracking Analyze (NTA) (Fig. 2F). Both EV subsets showed cup-shaped morphology under transmission electron microscopy (TEM), agreeing with previous reports on EV morphology^34^ (Fig. 2G and Fig. S10). Based on the guideline on EV analysis^35^, we performed western blotting (WB) on the expression of canonical EV marker proteins. CD63, CD9 and TSG-101, were detected in both NPs-carrying and NP-free EVs at the similar level (except TSG-101), while no expression of calnexin, a negative marker for EVs, was detected in both EV subsets (Fig. 2H). Similar result was obtained with R-AgNPs in HUVEC cells (Fig. S11). These results collectively demonstrate the vesicles we purified as EVs.

GW4869 is a compound that blocks ceramide-mediated inward budding of multivesicular bodies and thus prevents the release of EVs, especially exosomes^36^. After confirming that GW4689 treatment induced no or little cytotoxicity (Fig. 3A and Fig. S12), we used it to investigate the effect of inhibiting EV biogenesis on the intercellular exchange. We firstly checked whether GW4869 treatment could inhibit the production of EVs (both total EVs and NP-containing EVs). T-AgNP-containing donor cells were treated with GW4869, and the total EVs as well as NP-containing EVs were collected. The result of both particle number and the protein amount showed that GW4869 treatment indeed lowered the amount of total EVs (Fig. 3 B and C) and the fraction of NP-containing vesicles (Fig. 3 D and E). On the other hand, we found that the amount of freely released CPP-AgNPs was not affected by GW4869 treatment, indicating that GW4869 effect is specific to EV route (Fig. 3F). We then carried out the 3D intercellular exchange experiments. GW4869 treatment significantly lowered the intercellular exchange of T-AgNPs and R-AgNPs, and this result was observed with multiple donor/recipient pairs (Fig. 3 G and H, Fig. S13). Besides AgNPs, we found that the intercellular exchange of TAT-conjugated gold nanoparticles (T-AuNPs), and TAT-conjugated Dextran (T-Dextran), was also decreased by GW4869 treatment (Fig. 3 I and J). We further applied ionomycin, an ionophore that stimulate EV secretion^37,38^, to validate if the treatment could increase the intercellular exchange of CPP-NPs. Significant enhancement of the intercellular exchange of CPP-AgNPs in different donor/recipient pairs was observed after ionomycin treatment (Fig. S14A-D). Similar result was also obtained with T-AuNP and T-Dextran (Fig. S14 E and F). Overall, these results support the notion that CPP-NPs are exported inside EVs for intercellular exchange, and EV biogenesis is important for intercellular exchange of not only AgNPs, but also nonetchable metal NPs and organic polymers as well.

**Fig. 3.**
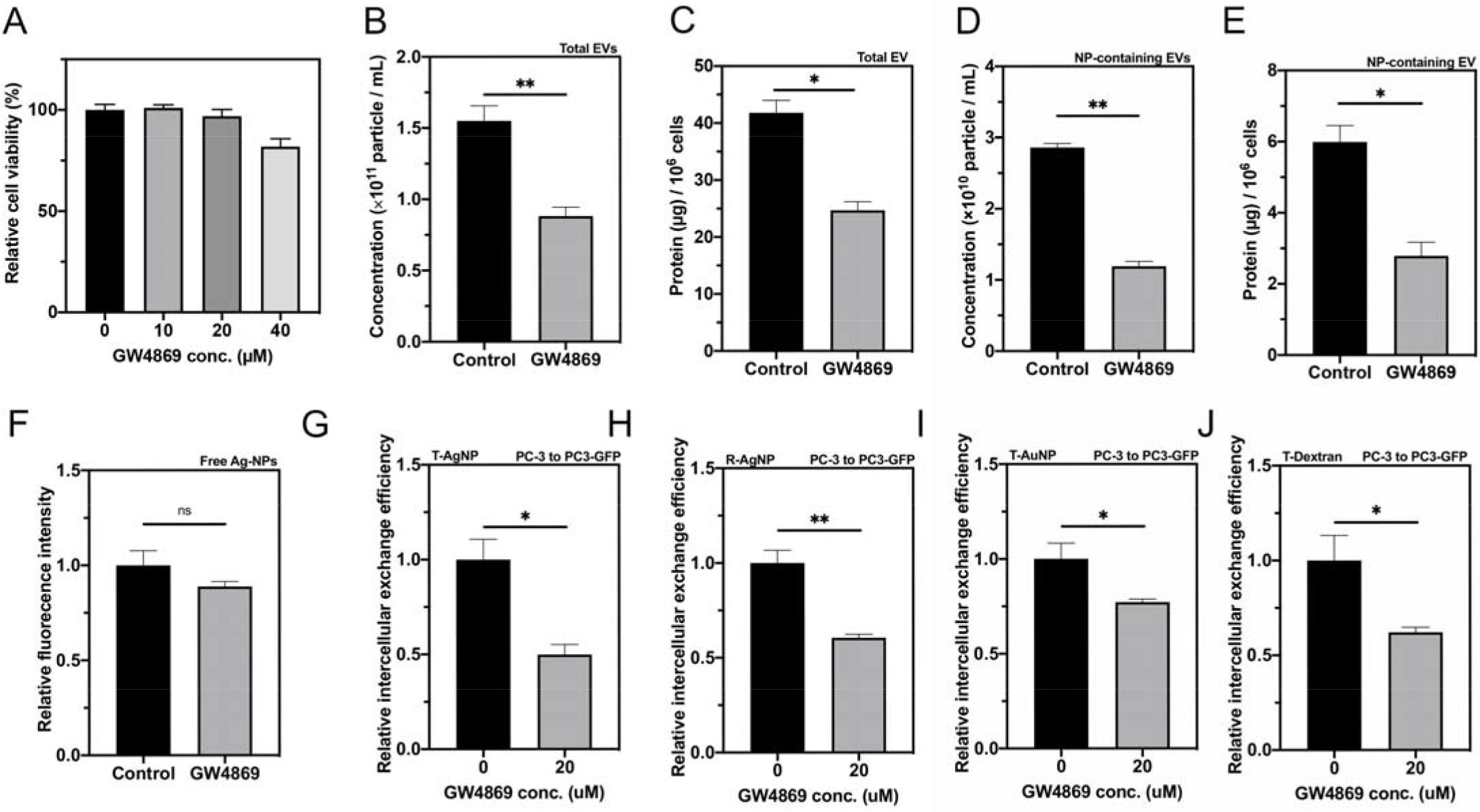
GW4869 treatment inhibited biogenesis of EVs and decreased the intercellular exchange of CPP-NPs. (A) Cell viability in the intercellular exchange assay with indicated concentration of GW4869 treatment (x axis) was tested and to that of without GW4869 treatment (y axis). (B)-(F) Exosome secretion inhibitor, GW4869, decreased secretion of total EVs and NP-containing EVs but not the release of free T-AgNPs. After feeding with T-AgNPs and washed with etchant, PC-3 cells were cultured DMEM (with EV free FBS) with 20 μM GW4869 for 48 h. After collected total EVs, NP-containing EVs and released free T-AgNPs as described in Methods, the particle concentration of total EVs (B) and NP-containing EVs (D) was quantified by NTA (y axis). The protein amount of total EV (C) and NP-containing EVs (E) was quantified by BCA assay (y axis). (F) The fluorescence intensity of released free T-AgNPs was measured as described in Methods and normalized to that of control group (y axis). (G and H) GW4869 decreased intercellular exchange of CPP-AgNPs. Intercellular exchange efficacy of T-AgNPs (G) and R-AgNPs (H) from PC-3 to PC3-GFP cells with 20 μM of GW4869 (x axis) was quantified as described in Methods and normalized to that of without GW4869 treatment (y axis). (I and J) GW4869 decreased intercellular exchange efficacy of T-AuNPs (I) and T-Dextran (J) from PC-3 to PC3-GFP cells with 20 μM of GW4869 (x axis) was quantified as described in Methods and normalized to that of without GW4869 treatment (y axis).Error bars indicate S.E.M., n=3. *P<0.05, **P<0.01 and ns, no significance (Student’s t-test).

### Re-entry of NP-containing EVs into recipient cells

Next, we evaluated the EV-mediated entry efficiency into recipient cells. EVs from a variety of donor cell types were isolated and labeled with a lipophilic dye. After normalization based on the particle numbers (Fig. S15A), they were incubated with recipient cells in both NP-free and NP-containing forms. Our results showed that loaded with NP or not, EVs enter the cells with similar efficiency (Fig. 4A). We also isolated NP-free EVs from different types of parent cells and tested their cell entry efficiency in a variety of cell types. The cell entry efficiency of EVs varied, which depends on both parent and recipient cell types (Fig. S15B-D). Second, we verified the transport rate of EVs in the collagen matrix. Both NP-free and NP-containing EVs seemed to rapidly diffuse through the collagen and colocalized with PC3-GFP cells at the bottom of collagen within 30 minutes (Fig. S15E). The 3D reconstruction of confocal images of recipient cells confirmed that the intact labeled EVs were taken up and stayed inside recipient cells (Fig. S15F). This result indicates that collagen gap and matrix allow efficient transport and exchange of EVs.

**Fig. 4.**
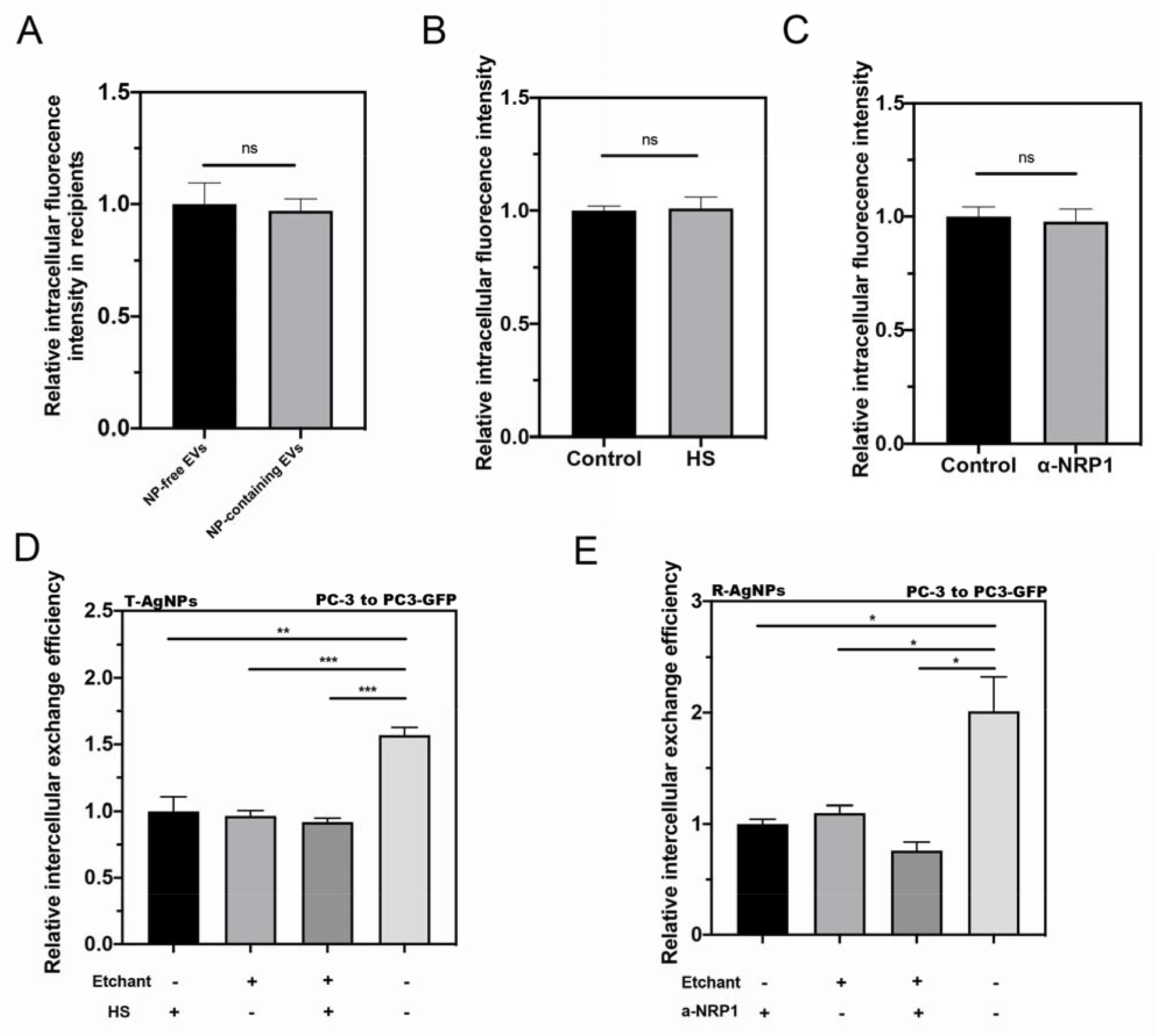
Re-entry of NP-containing EVs into recipient cells. (A) Similar cell entry efficiency of NP-free and NP-containing EVs in recipient cells. NP-free EVs and NP-containing EVs were isolated from T-AgNPs engulfed PC-3 cells and labeled with Dil. After incubating with PC-3 recipient cells for 2 h, the fluorescence signal of EVs inside the cells was detected by confocal microscopy, quantified by Image J, and normalized to that of NP-free EVs (y axis). (B and C) Similar cell entry efficiency of Dil labeled EVs in recipient cells under treatment of HS (B) and «-NRP1 (C). Total EVs were isolated from PC-3 cells and labeled with Dil. After incubating with PC-3 cells for 2 h with indicated treatment (x axis), the intracellular fluorescence intensity was measured and normalized to control groups (y axis). (D and E) Re-entry of EV-carrying CPP-AgNPs did not rely on ligand-receptor interaction. (D) Intercellular exchange efficacy of T-AgNPs from PC-3 to PC3-GFP with indicated treatments (x axis) was quantified as described in Methods and normalized to that of HS treated only group (y axis). (E) Intercellular exchange efficacy of R-AgNPs from PC-3 to PC3-GFP with indicated treatments (x axis) was quantified as described in Methods and normalized to that of «-NRP1 treated only group (y axis). Error bars indicate S.E.M., n=3. *P<0.05, **P<0.01, *** P<0.001 and ns, no significance (Student’s t-test).

Lastly, the endocytosis of T-AgNPs and R-AgNPs into cells are mediated by the interaction with their receptors, heparin sulfate (HS) proteoglycans and neuropilin-1 (NRP1), respectively^26^. Here, we set out to test whether freely released CPP-NPs and NP-containing EVs enter the recipient cells in the same manner. We first confirmed that the uptake efficiency of EVs was not affected by HS and NRP1 antibody (Fig. 4B and 4C). Using soluble HS and NRP1-blocking antibody, we were able to reduce the intercellular exchange of T-AgNPs and R-AgNPs, respectively. This was performed without etching, and the levels of reduction were very similar to that of etching treatment. In the presence of constant etching, these cell entry inhibitors exhibited little effect (Fig. 4 D and E). These results demonstrate that these inhibitors can effectively block the re-entry of freely released CPP-NPs into recipient cells, while NP-containing EVs are resistant to them. These data support the notion that while freely released CPP-NPs still rely on the same ligand-receptor interaction for cell entry, NP-containing EVs explore a distinct pathway to re-enter the recipient cells.

### EV biogenesis is important for NP delivery into solid tumors *in vivo*

Last, we set out to validate the importance of EV biogenesis in NP delivery *in vivo*. Here, we used iRGD to functionalize AgNPs, as it was shown that iRGD can induce the vascular and tumor penetration of a variety of NPs, including AgNPs, *in vivo*^20,39,40^. To inhibit EV biogenesis, we intratumorally injected GW4869 into mice bearing 4T1 breast tumor. By using the TUNEL staining of tumor tissues, we confirmed that GW4869 showed little toxicity to tumor cells (Fig. S16). The homing study by intravenous injection of iRGD-AgNPs showed that compared to control group, GW4869 treatment significantly reduced the overall accumulation of iRGD-AgNPs in the tumor tissue (Fig. 5A and Fig. S17A). We also quantified the penetration distance of iRGD-AgNPs from the nearest blood vessel (Fig. S18), and found that GW4869 treatment significantly decreased the penetration distance of iRGD-AgNPs in tumor tissue (Fig 5B). We validated this observation with an orthotopic pancreatic ductal adenocarcinoma tumor model (Fig. 5C, 5D and Fig. S17B). Notably, GW4869 was injected intravenously in this study, to verify whether its effect remains after systemic administration. A significant reduction of iRGD-AgNP accumulation was also seen in tumor tissue after GW4869 treatment, as well as the vascular penetration distance.

**Fig. 5.**
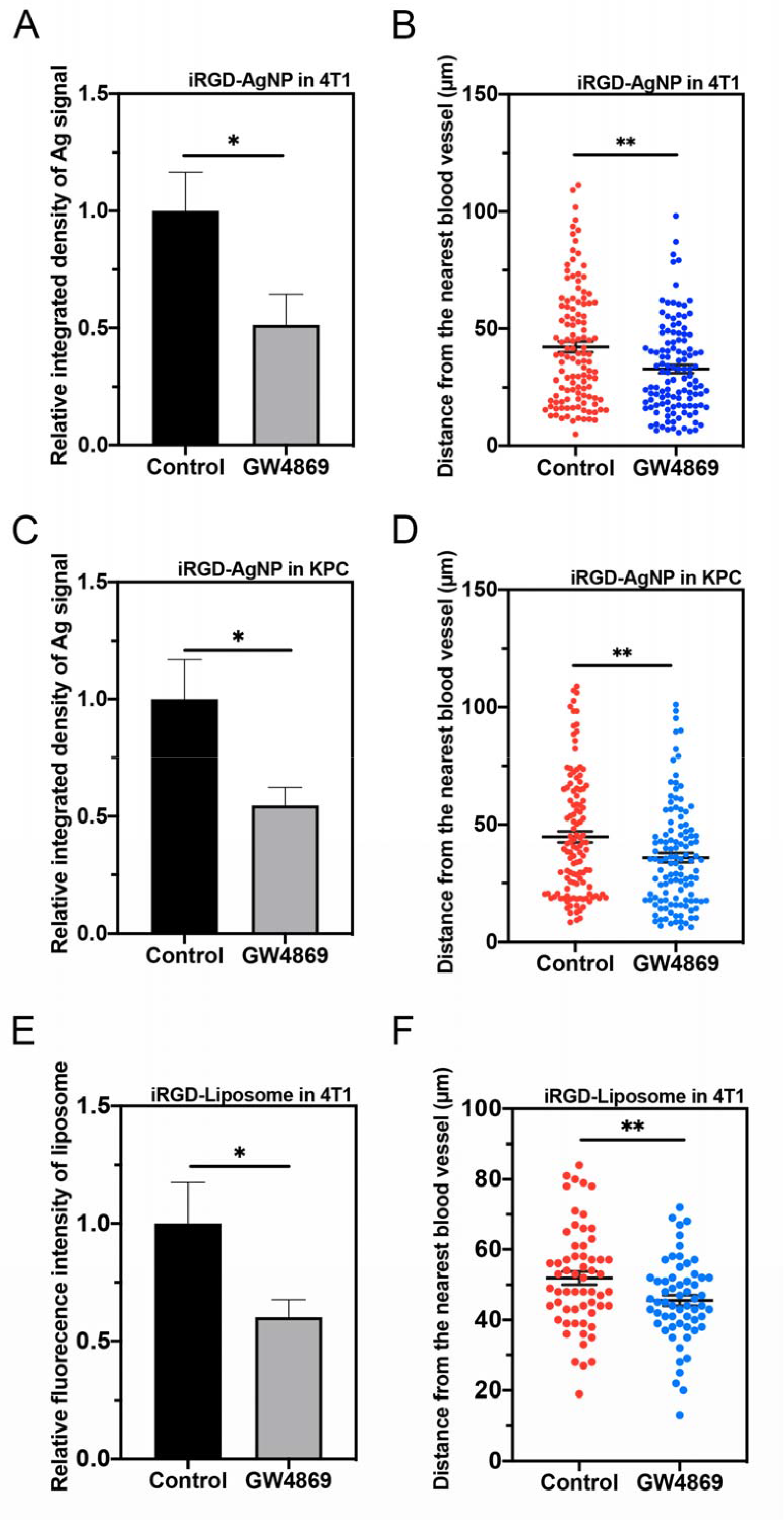
*In vivo* demonstration of EV importance in iRGD-NP delivery into solid tumors. GW4869 decreased the accumulation and penetration of iRGD-NPs in 4T1 and orthotopic pancreatic ductal adenocarcinoma tumor model (indicated as KPC in the figure) *in vivo*. 4T1 tumor bearing mice received 20 μL of GW4869 (40 μM) via intratumoral injection each day for 5 days. Orthotopic pancreatic ductal adenocarcinoma tumor bearing mice received GW4869 at a dosage of 2.5 μg/kg body weight via intraperitoneal injection every other day for 5 injections. 24 h after the last injection, 50 μL of iRGD-AgNPs (O.D 40) or 100 μL of iRGD-Liposome was intravenously injected and circulated for 4 h. Tumor was excised and sectioned for AgNP and blood vessel detection as described in Methods. (A) Semi-quantitative analysis of Ag signal intensity in 4T1 tumor tissue by ImageJ, and normalized to that of control group (y axis). (B) Quantitative analysis of distance of Ag signal to the nearest blood vessel in 4T1 tumor by ImageJ. (C) Semi-quantitative analysis of Ag signal intensity in KPC tumor tissue by ImageJ, and normalized to that of control group (y axis). (D) Quantitative analysis of distance of Ag signal to the nearest blood vessel in 4T1 tumor by ImageJ. (E) Semi-quantitative analysis of iRGD-Liposome signal intensity in 4T1 tumor tissue by ImageJ, and normalized to that of control group (y axis). (F) Quantitative analysis of distance of iRGD-Liposome to the nearest blood vessel in 4T1 tumor by ImageJ. The *in vivo* experiment for iRGD-AgNP was performed in 3 mice per group. 5 images from each tumor tissue were used to analyze Ag signal intensity in each group. 40 sliver signal from each tumor tissue were applied to analyze the penetration distance. The *in vivo* experiment for iRGD-Liposome was performed in 2 mice per group. 3 images from each tumor tissue were used to analyze fluorescent signal intensity in each group. 10 liposome signal from each image were applied to analyze the penetration distance. Error bars indicate S.E.M.. *P<0.05, **P<0.01 (Student’s t-test).

Next, we set out to study whether GW4869 treatment could also affect the accumulation and penetration of organic NPs. Liposome has been well studied and widely used as drug delivery system^9,41^. We prepared iRGD conjugated liposome (iRGD-Liposome) to study the effect of GW4869 treatment on tumor accumulation and penetration. Similar to the experiment of iRGD-AgNPs, GW4869 was intratumorally injected into mice bearing 4T1 breast tumor to inhibit biogenesis of EVs, followed by intravenous injection of iRGD-Liposome for homing. The results showed that GW4869 treatment significantly reduced the overall tumor accumulation and vascular penetration distance of iRGD-Liposome (Fig. 5 E and F). Together, these data suggest that EV biogenesis is important for extravasation, and more importantly, the deeper penetration of iRGD-NPs (both inorganic and organic) into the extravascular regions through the transcellular route.

## Discussion

Here we provide a 3D assay to study an important and yet understudied part of transcellular transport for NPs, intercellular exchange. Aided by CPPs and etching technology, this assay is easy to establish, better mimics the physiological environment for cells, and is able to distinguish different intercellular exchange routes. Our results prove that a significant fraction of NPs are exported inside EVs, which carry NPs into recipient cells independent of the original cell-penetrating ligands on them. Our study unveils a novel role of EVs in the transcellular transport of NPs.

Despite its importance, transcellular transport process is poorly characterized for NPs and other types of macromolecule payloads. It consists of multiple interconnected steps, which is too complicated to investigate under *in vivo* conditions. *In vitro*, the endocytic machineries in general, and the cell entry and intracellular transport of NPs, are much better understood than exocytosis and intercellular material exchange^17,18,42^. Therefore, there is great need to study the intercellular exchange, not only for NP applications but also for a better understanding of exocytosis and intercellular communication in general. Years of studies on transcytosis have shed light in cargo transport across one layer of cells, and generated useful cellular assays^43–45^. However, it differs from intercellular exchange in that transcytosis includes the initial endocytosis into donor cells, while excludes the re-entry into recipient cells. The re-entry step is crucial, in our opinion, as NPs and other payloads need to at least enter one more layer of cells after crossing the endothelium for their therapeutic effect. This is also an important cell biology question in regard to how exocytosed materials interact with cells. Furthermore, transcytosis assays (e.g. Transwell assay) measures the entire process from the entry into the first cell layer to re-entry into recipient cells, while it is difficult to ensure no leakage in the intercellular gaps for NPs of various sizes and shapes. To solely focus on the intercellular exchange, we thus developed the described 3D assay.

The key to our intercellular exchange 3D assay is etching. After CPP-AgNP internalization, etching ensures that no extracellular or cell surface bound AgNPs exist. In this way, all AgNPs in recipient cells can only come from the export from donor cells. The etchant is a small molecule compound that can easily access all extracellular spaces, acts very rapidly (~seconds *in vitro*) and effectively, and is nontoxic to cells^20^. Therefore, constant etching can dissolve any freely released AgNPs from donor cells before they can reach the recipient. Our results also confirmed that etching process barely affected the ability of secretion / uptake of EVs by donor / recipient cells, respectively. Together, etching can help us ensure that we are observing the intercellular exchange events and help distinguish the direct release route from others. Besides etching, our assay has other advantages. Bulk processing of donor cell uptake helps eliminate the interference from the cell entry into donor cells and their subcellular transport status. The collagen gap, meanwhile, is an easy and effective way to prevent direct cell-cell contact. Together, these properties ensure that we are exclusively and precisely monitor and quantify the intercellular exchange events and enable us to differentiate different transfer routes. Our results demonstrated that CPP-NPs are exported either freely or enclosed inside EVs, but not exchanged via direct cell-cell contact. It is very difficult, if not possible, to distinguish these routes *in vivo* or with existing cellular assays. Besides AgNPs, we also demonstrated that EVs are important for intercellular exchange of nonetchable NP type (AuNPs and Dextran), supporting the generality of our findings.

The use of CPPs, besides aiding in the cell entry of NPs, help elicit the impact of ligand-receptor interactions on intercellular exchange. AgNPs are versatile in changing their physicochemical characteristics (e.g. sizes, shapes, surface charges), and have been shown to be coupled with a wide variety of cell-penetrating ligands beyond CPPs^46–48^. Our results showed that the intercellular exchange efficiency varies with CPP types and AgNP sizes. Further investigations in this regard will elucidate the impact of various NP properties and ligand functionalization on their intercellular exchange efficiency. This assay format is also flexible to study different donor/recipient cell types, matrix compositions and environmental conditions. For example, we investigated in this study the intercellular exchange between tumor cells (PC-3 / PC3-GFP), endothelial cells and tumor cells (HUVEC / PC3-GFP), tumor cells and fibroblasts (4T1 / NIH-3T3), immune cells to tumor cells (THP-1 / PC3-GFP). Finally, the procedure of this 3D assay is simple enough that we can envision to adapt it for high throughput screening. Such screens may quickly identify genetic factors or chemical compounds that can up- or down-regulate the intercellular exchange, and likely the tissue penetration/delivery, of NPs.

We have set up this assay to answer a fundamental cell biology question: how NPs are exported from one cell and transferred into another. In the meantime, we tried to simulate the *in vivo* scenario. NP extravasation has been the central focus for its delivery into solid tumors and likely other solid tissues. Therefore, we used HUVEC, a widely used endothelial cell line, as the donor cells. Plus, extravasation is only the first step for NP delivery, and it is of great scientific and therapeutic significance to understand the intercellular material exchange beyond the vasculatures. Thus, a wide variety of cell types were also tested here, including tumor cells (PC-3 and 4T1), fibroblasts (3T3) and immune cells (THP-1). A variation of intercellular exchange efficiency was seen with different cell types in our results, agreeing with previous reports that EV biogenesis and cell uptake are highly cell type specific^49–51^. More thorough investigations are needed to elicit the underlying mechanism.

EVs have long been recognized as an important pathway for intercellular communications by exchanging various types of payloads between cells^31,32,52,53^. EVs such as exosomes are also actively being developed as the carriers for drug delivery^54–56^. On the other hand, while cellular transfer of NPs and their payloads has received increasing attention^57–59^, the role of EVs in this process remains to be elusive. Here, our major conclusion is that EVs mediate a significant part of intercellular exchange of CPP-NPs. We proved that CPP-NPs can be exported inside EVs, and EVs can deliver themselves and the enclosed NPs into another cell. Using GW4869, we demonstrated that EV biogenesis is important for NP intercellular exchange *in vitro*, and tissue penetration *in vivo*. The *in vivo* result is particularly intriguing. To our best knowledge, this is the first *in vivo* evidence implicating the involvement of naturally occurring EVs in the delivery of both organic (liposome) and inorganic (AgNP) NPs into solid tissues (e.g. tumors), and their tissue penetration beyond the vasculatures. Inorganic NPs, such as AgNPs, can remain intact at least up to 24-h after cell entry^20^, so it is reasonable to conclude that intact NPs are packaged into EVs. On the other hand, studies have reported that liposomes dissemble after entering cells and released their cargo^60–62^. While it is challenging to fully distinguish intact and dissembled liposomes, especially *in vivo*, evidence from our group and others suggests that at least a fraction of liposomes may be still intact during EV-mediated intercellular exchange. It has been showed that intact liposomes can be taken up by cells^63^ and the formulation of liposomes decides their stability inside cells^64^. The cholesterol used in our liposomes stabilized the overall structure and prolonged their half-life inside cells^64^. We recently showed that our liposomes remain largely intact in the tumor tissue up to a few hours after intravenous injection^65^. Further studies are needed to accurately quantify the ratio of intact organic NPs that can be exocytosed inside EVs.

Lastly, we found that EV-enclosed NPs do not require coupled CPPs for cell entry after the initial uptake by donor cells. It is reasonable to speculate that as NPs (as well as other cargo) are fully encapsulated and thus their specificity towards recipient cells likely depends on EVs themselves. The cell uptake specificity of EVs mainly relies on their surface proteins and their interactions with the receptors on recipient cells, which are again highly variable and depend on the donor and recipient cell types, respectively^51,52^. Our study supports this notion in that the cell entry efficiency of EVs varies with parent and recipient cell types, and NP-containing EVs show no or little difference from NP-free ones. Overall, our results call for more careful design of ligand-functionalized NPs when aiming at solid organs/tissues like tumors, and highlights the importance of EV biology (surface proteomics, transport dynamics and cell uptake specificity) in improving the tissue penetration and therapeutic efficacy of NPs.

## Materials and methods

### Cell Lines and Cell Culture

Human prostate cancer cell line PC-3, human monocyte cell line THP-1 and mouse breast cancer cell line 4T1 were purchased from American Type Culture Collection (ATCC CRL-1435, VA, USA). PC3-GFP and KPC cells was a gift of Dr. Erkki Ruoslahti’s lab. Primary Human Umbilical Vein Endothelial Cells (HUVECs) and mCherry-labeled mouse fibroblast cell line NIH-3T3 were gifts from Dr. David K. Wood, University of Minnesota. PC-3, 4T1, PC3-GFP KPC and NIH-3T3 cells were cultured in Dulbecco’s modified Eagle’s medium (DMEM, cat. no. 16777-129, VWR international, LLC.) supplemented with 10% fetal bovine serum (FBS, cat. no. 35-011-CV, Corning), and 1% penicillin−streptomycin (10000 U/mL) (cat. no. SV30010, Thermo Fisher Scientific Inc.). HUVECs were cultured in Endothelial Cell Growth Medium-2 BulletKit (EGM-2, cat. no. CC-3162, Lonza Inc., ME, USA). THP-1 cells were cultured in RPMI-1640 medium. All cells were maintained in a 37 °C humidified incubator with 5% CO_2_. For extracellular vesicle (EV) isolation, cells were cultured in DMEM with EV-free FBS.

### Preparation of nanoparticles

Detailed preparation method of nanoparticles used in this study can be found in supplementary information.

### Preparation of the etchant

20x Etchant stock for *in vitro* use:

Reagent A: Tripotassium hexacyanoferrate (III) (K_3_Fe(CN)_6_, Sigma, CAS# 13746-66-2) was typically dissolved in DPBS (Hyclone, cat. no. SH30028.02) at 0.20 M and stored in the dark.
Reagent B: Sodium thiosulfate pentahydrate (Na_2_S_2_O_3_ : 5H_2_O, Sigma, CAS# 10102-17-7) was typically dissolved in DPBS at 0.2 M.
These stocks were stored at room temperature in 50 mL polypropylene tubes in the dark for at least a month without issue. When used for *in vitro* study, Reagent A and B were freshly mixed at 1:1 (V/V) and diluted 20x by medium.

### Establishment of intercellular exchange assay

#### Preparation of donor cells

Donor cells were cultured in 10 cm dishes or 6-well plates. When cells reached 70-80% confluency, the culture medium was replaced with nanoparticle-contained medium and incubated at 37 °C for 4 h. After incubation with nanoparticles, 20 μL of etchant was added into each well and was rocked gently for 30 s. Then the etchant was removed and the cells were washed 3 times with PBS. Cells were trypsinized with 0.05% Trypsin-EDTA (cat. no. 17-161E, Thermo Fisher Scientific Inc.). Cells were counted and resuspended with medium for further use.

#### Preparation of recipient cells

Recipient cells were trypsinized with 0.05% Trypsin-EDTA when cells reached 70-80% confluency. Cells were counted and resuspended with medium for further use.

#### Preparation of intercellular exchange assay (All operations were carried out on ice)

The detailed optimization process for conditions of intercellular exchange assay is provided in the supplementary information. The optimal condition of the intercellular exchange assay is listed below, if not otherwise indicated: Type I collagen (Collagen Type I, Rat tail high concentration, 8.91 mg/mL, ref. no. 354249, Corning) was mixed with sterile 10x PBS, 1N NaOH, H_2_O and 9×10^4^ recipient cells in 200 μL cell culture medium, making the final concentration of collagen to 2 mg/mL and pH around 7.4. Then 30 μL of recipient cell-collagen mixture was firstly added into each well in a 96-well plate. The plate was then placed into the cell incubator (37 °C) for 15 min to allow for collagen polymerization. An acellular collagen gap mixture was prepared using the procedure above without the addition of cell suspension. After the recipient layer polymerized, 30 μL of 2 mg/mL collagen gap mixture was added into each well and incubated at 37 °C for another 15 min. Then, 1.8 ×10^5^ donor cells in 200 μL medium with etchant was added on top of collagen gap and the plate was moved into the incubator for 24 h. After the intercellular exchange was completed, 100 μL of medium in each well was removed from the top and 100 μL of 1% collagenase (Sigma, cat. no. C9263-1G) in FBS free medium was added into the well and incubated at 37 °C for 20 min. The mixture was then transferred into Eppendorf tubes and centrifuged at 300 RCF at 4 °C for 10 min. Finally, the cell pellet was fixed with 4% formalin (Sigma, cat. no. HT501128-4L) and stored at 4 °C. The intercellular exchange efficacy between donor and recipient cells was measured by flow cytometry using a BD FACS Calibur flow cytometer (BD Biosciences, San Jose, CA). The intercellular exchange efficacy was calculated as:

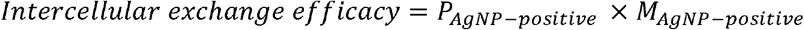

where *P*_*AgNP-positive*_ stands for percentage of AgNP-positive recipient cells and *M*_*AgNP-positive*_ stands for mean fluorescence intensity in AgNP-positive recipient cells.

To prepare donor cells in collagen format, the same procedure was applied as preparing recipient-collagen mixture except replacing recipient cells with donor cells. After recipient layer and gap layer were polymerized, 30 μL of donor layer mixture was added on top of the collagen gap and incubated at 37 °C for 15 min to polymerize. Then 200 μL of medium was added into each well. Other conditions were the same as described above.

To perform intercellular exchange assay with different gap concentrations, the final collagen concentration was adjusted with sterile 10x PBS, 1N NaOH and H_2_O to make the final concentration of collagen to 2, 4 and 8 mg/mL and a pH of approximately 7.4. Other conditions were the same as described above.

For groups without gap or etchant, after the recipient layer was polymerized, 200 μL of donor cells suspension was seeded directly onto the recipient cell-collagen layer and incubated for 24 h. Other conditions were the same as described above.

### Intercellular exchange with GW4689 and ionomycin

To study the effects of GW4689 (Sigma, cat. no. D1692-5mg) on intercellular exchange, 10, 20 and 40 μM of GW4689 was added into the medium under constant etching, respectively. After 24 h of incubation, cells were collected using the same method described above and analyzed by flow cytometry.

To study the effects of ionomycin (Sigma, cat. no. I3909-1mL) on intercellular exchange, after incubating with nanoparticle for 4 h and being washed with etchant and PBS for 3 times, donor cells were incubated with 1.25 μM of ionomycin (in complete DMEM medium) at 37 °C for 10 min. After that, donor cells were washed with PBS for 3 times and the intercellular exchange assay was carried out using the same method described above and analyzed by flow cytometry.

### EV Isolation and density gradient ultracentrifugation

Conditioned medium was collected from donor cell cultures after 48 h of incubation in FBS-free medium. EVs were isolated using ultrafiltration method as described previously^34^. Briefly, conditioned medium was harvested and centrifuged at 4 °C, 300x g for 10 min followed by 2000x g for 10 min to remove cells and debris. Then, the supernatant was transferred to a Centricon Plus-70 Centrifugal Filter (Sigma, UFC710008) and centrifuged at 4 °C, 3500x g for 40 min. The concentrated medium was collected using reverse spin at 1000x g for 2 min. Density gradient ultracentrifugation were carried out as described previously^33^. Briefly, a series of gradient solution (40 % (w/v), 20 % (w/v), 10 % (w/v) and 5 % (w/v) solutions of iodixanol) were prepared by diluting a stock solution of OptiPrep™ (60 % (w/v) Sigma, cat. no. D1556-250ML) with 0.25 M sucrose, 10 mM Tris–HCl, pH 7.5 solution. The isolated EVs were suspended in 0.5 mL of 5% gradient and then layered on top of a gradient consisting of 10%, 20%, and 40% OptiPrep (3 mL for each gradient). Gradients were centrifuged using a SW 40 Ti rotor at 100000g for 18 h at 4 °C. Fractions of 1 mL were collected from the top of the gradient. NP-free fraction (fraction 7&8) and NP-containing fraction (fraction 10 & 11) were diluted in PBS (1:25) and centrifuged at 100000g for 90 min at 4 °C. The pellets were resuspended in cold PBS and were stored at 4 °C for further use.

### Transmission electron microscopy (TEM) imaging and Dynamic light scattering (DLS) analysis

EVs were visualized using TEM according to Lobb et al^34^. Briefly, 2 μL of exosome suspension was fixed in 50 μL of 2% paraformaldehyde. 2 μL of this mixture was transferred onto each of 2 Formvar-carbon coated electron microscopy grids. Membranes were covered for 15 minutes. A 100 μL drop of PBS was placed on a sheet of parafilm and grids transferred with the sample membrane side facing down using clean forceps for 2 minutes. The grids were transferred to a 50 μL drop of 1% glutaraldehyde for 5 minutes before transferring to a 100 μL drop of distilled water for 2 minutes. This was repeated 7 times for a total of 8 water washes. To contrast the samples, grids were transferred to a 50 μL drop of uranyl-oxalate solution, pH 7, for 5 minutes before transferring to a 50 μL drop of methyl-cellulose-UA (a mixture of 4% uranyl acetate and 2% methyl cellulose in a ratio of 100 μL/900 μL, respectively) for 10 minutes, placing the grids on a glass dish covered with parafilm on ice. The grids were removed with stainless steel loops and excess fluid blotted gently on Whatman no.1 filter paper. Grids were left to dry and stored in appropriate grid storage boxes. Grids were observed with JEM 1011 tranmssion electron microscope at 90 kV.

For DLS measurement, EVs were diluted to 500 μL with cold PBS and the diameter of EVs were measured using PAN185 DLS Analyzer.

### Confirmation of AgNPs in isolated EVs

Total EVs isolated from donor cell culture medium or NP-containing EVs isolated by density gradient ultracentrifuge were washed with PBS and re-suspended in 50 μL of cold PBS. Then samples were aliquoted into 3 x 50 μL. The three aliquots were treated with 50 μL of cold PBS, 50 μL of etchant and 50 μL etchant with 0.2% Triton X-100 (1:1), respectively. The 4 mixtures were then transferred to a black 96-well plate. The fluorescence intensity of CF647 was measured within 10 min using a SpectraMax M2 plate reader (Molecular Devices, Inc.).

### Western blot analysis

EV samples were lysed with lysis buffer (Cell Signaling Technology, Danvers, MA) supplemented with phenylmethylsulfonyl fluoride protease inhibitor on an ice bath for 30 min. Following centrifugation of the lysates at 14000xg and 4°C for 20 min, the supernatant was collected for Western blot (30 μg of total protein/lane). Protein concentration was measured using the Pierce™ BCA^®^ Protein Assay Kits (Thermo Fisher Scientific Inc., cat.no. 23227). CD63 antibody (1:500 Thermo Fisher Scientific Inc., cat.no. 10628D), CD9 antibody (1:500 Thermo Fisher Scientific Inc., cat.no. 10626D), TSG-101 antibody (1:500 Thermo Fisher Scientific Inc., cat.no. MA5-32463) and calnexin antibody (1:500 Thermo Fisher Scientific Inc., cat.no. 10427-2-AP) were used for immunoblotting.

### GW4689 inhibitory effect on EVs secretion and exocytosis of T-AgNPs

When PC-3 cells reached 80% confluency in 15 cm culture dish, the culture medium was replaced with nanoparticle-containing medium and incubated for 4 h. After incubation with nanoparticles, cells were washed with etchant and PBS and then cultured in DMEM with 40 μM of GW4869 for 48 h. Total EVs, NP-free EVs and NP-containing EVs were then isolated as described above. Medium was collected for analysis of the exocytosis of T-AgNPs. The size distribution and particle concentration of different EVs were quantified by NTA. For analysis of exocytosis of T-AgNPs, collected medium was concentrated with centrifugal filter tube with 10,000 MWCO into 1mL. Then Bioworld Heparin-Coated Plate (Thermo Fisher Scientific Inc., cat.no. 50197531) was used to capture released T-AgNPs in the concentrated medium. After washing with PBS for 3 times, 150 μL of PBS was added into each well and the fluorescence intensity of CF647 was measured using a SpectraMax M2 plate reader (Molecular Devices, Inc.).

### Re-entry of EVs and EV penetration in collagen matrix

All EVs were labeled with Dil (Thermo Fisher Scientific Inc., cat.no. D3911). Briefly, 2 μL of Dil working solution (2 mg/mL) was added into 100 μL of EVs solution. The mixture was mixed well with pipet and incubated at 37°C for 30 min. After incubation, the mixture was centrifuged at 3000g for 10 min. The supernatant was collected and EVs were normalized to the same protein amount for further use.

For studying re-entry of EVs into cells, EVs from PC-3, 4T1 and HUVEC cells were collected and labeled as described above and normalized to the same protein amount. PC-3, 4T1 and HUVEC cells were cultured in 6-well plate and incubated with labeled EVs from different origins for 4 h. After washing with PBS for 3 times, cells were incubated with Hoechst 33342 (Thermo Fisher Scientific Inc., cat.no. 62249) for 15 min and washed with PBS. Fluorescence images were taken using EVOS M5000 (Thermo Fisher Scientific) microscope and the fluorescence intensity of Dil was analyzed by ImageJ.

To study the penetration of EVs in collagen matrix, a single layer of PC3-GFP cells were seeded in 8 Chamber Polystyrene Vessel Tissue Culture Treated Glass Slide (Falcon, cat. no. 354108) and cultured overnight. After cells attached to the slides, medium in chambers was discarded and the collagen mixture prepared as described above was added on top of cells and incubated at 37 °C for 15 min to polymerize. Then Dil labeled NP-free EVs and NP-containing EVs were added into the chamber with 300 μL medium (with EV-free FBS) and incubated at 37°C for 30 min. Medium was removed and the chamber was washed with PBS. Cells were fixed with 300 μL 10% formalin for 4 h at room temperature. Formalin was removed and 800 μL O.C.T. compound (Tissue-Tek, SAKURA^®◻^, cat. no. 4583) was added into the chamber. Then the whole chamber was transferred into −80 °C overnight. The next day, the frozen bulk gel was taken out of the chamber and frozen sections of bulk gel was prepared according to standard cryo-section protocol. Fluorescence images were taken using EVOS M5000 microscope (Thermo Fisher Scientific).

### In vivo study

All animal studies were carried out in compliance with the National Institutes of Health guidelines and an approved protocol from University of Minnesota Animal Care and Use Committee. The animals were housed in a specific pathogen-free facility with free access to food and water at the Research Animal Resources (RAR) facility of the University of Minnesota.

To establish 4T1 orthotopic tumor model, 100 μL of 1×10^7^ cells/mL 4T1 cells suspended in PBS were injected into the mammary fat pad of female Balb/c mice. Once the average tumor volume reached 80 mm^3^, mice in the control group were intratumorally injected with 20 μL of 10% DMSO in PBS daily for 5 days. Mice in GW4869 group were intratumorally injected with 20 μL of GW4869 (40 uM final conc.) daily for 5 days.

The orthotopic pancreatic ductal adenocarcinoma tumor model was established according to a reported protocol^66^. Briefly, surgery procedure was performed to expose the entire pancreatic body together with spleen to the outside of the peritoneal cavity. 100 μL of 1×10^7^ cells/mL KPC cell-Matrigel (ref. no. 354234, Corning) mixture was injected into the tail of the pancreas. For orthotopic pancreatic ductal adenocarcinoma tumor, 14 days post-surgery, mice in control group were intraperitoneally injected with 200 μL of 10% DMSO in PBS every other day for 5 times. Mice in GW4869 group were intraperitoneally injected with 200 μL of GW4869 (2.5 μg/kg body weight) every other day for 5 times.

24 h after the last GW4869 injection, 50 μL of iRGD-AgNPs (O.D 40) / 100 μL of iRGD-Liposome / 50 μL of AgNPs (O.D 40) / 100 μL of Liposome was intravenously injected and circulated for 4 h. Animals were then anesthetized with Avertin and underwent heart perfusion before tumor tissue excision.

To stain blood vessels for quantifying penetration distance, after rehydrating with PBS, tissue section was covered with 200 μL of PBS containing 1% BSA and 0.1% Triton X-100 (blocking solution) incubated at room temperature for 1 h. The sections were washed three times with PBS and then incubated with primary antibody anti-CD31 (Rat Anti-Mouse, Thermo Fisher Scientific Inc., cat. no. MA1-40074) with a 1:200 dilution in blocking buffer at 4 °C overnight, followed by secondary antibody (Peroxidase-AffiniPure F(ab’)2 Fragment Goat Anti-Rat IgG (H+L), Thermo Fisher Scientific Inc., cat. no. NC9810135) diluted (1:200) in blocking buffer at RT for 1 h. AEC Substrate Kit (Thermo Fisher Scientific Inc., cat. no. NC9821752) was then used to stain blood vessels in tumor tissue.

Liposome in tumor tissue were stained with primary anti-FITC antibody (Invitrogen, A889) in blocking buffer at 4 °C overnight, followed by secondary antibody (Donkey Anti-Rabbit IgG H&L (Alexa Fluor® 488), Thermo Fisher Scientific Inc., cat. no. A-31572) diluted (1:200) in blocking buffer at RT for 1 h. After washing with PBS, sections were mounted in DAPI-containing mounting medium (Vector Laboratories, Burlingame, CA) with a coverslip and examined under fluorescence microscope EVOS M5000 (Thermo Fisher Scientific). AgNPs in tumor tissue were stained with LI Molecular Probes Silver Enhancement kit (Thermo Fisher Scientific Inc., cat. no. 24919) as previously described^20^. Briefly, collected tumor tissue was washed in PBS and “snap frozen” in liquid nitrogen. Cryosections of 7 μm thick were prepared and fixed in 4% PFA. After washing in PBS, the sections were incubated with etchant for 10 s. Then slides were washed in PBS twice followed by two washes in water. Tissues was blocked for 20 min with a blocking solution of glycine 50 mM pH 7.8, 0.04% sodium azide, 0.4% Triton-X 100 and 2% citrate sodium. The LI Molecular Probes Silver Enhancement kit was applied to stain silver for 30 min with fresh solution added every 10 min. After that, samples were washed in water for 3 times and coverslips were applied with cytoseal mount media.

The whole tumor scan images were taken using Huron TissueScope LE (Huron Technologies International Inc.). Other images were taken using EVOS M5000 (Thermo Fisher Scientific) microscope. The integrated density of Ag signal and penetration distance were analyzed by ImageJ.

### Statistical Analyses

All quantified data are presented as mean ± S.E.M. (standard error of the mean). All statistical analyses were performed using the GraphPad Prism software. Statistical significance was considered at P values lower than 0.05. P values are shown as *P ≤ 0.05, **P ≤ 0.01, ***P ≤ 0.001, and ****P ≤ 0.0001. No outliers were excluded in this study. The methods of statistical analyses have been indicated in figure legends. All comparisons between two experimental groups were performed by unpaired two-tailed Student’s t test.

## Author Contributions

H-B.P. designed the project. T.T., K.A.C. and D.K.W carried out collagen microtissue synthesis and experiment. X. W. and Y. W. carried out the rest of the study. X.W. and H-B.P. wrote the manuscript.

## Acknowledgements

Research reported in this publication was supported by grants from the from the National Institute of Health (R01CA214550, R01GM133885, R21EB022652) and the State of Minnesota (MNP#19.08). The content is solely the responsibility of the authors and does not necessarily represent the official views of the National Institutes of Health.

## Conflict of Interest

H-B.P. is a shareholder of Cend Therapeutics, Inc.

## Notes

### Competing Interest Statement

The authors have declared no competing interest.

### Summary of Updates

Manuscript and supplemental files updated.

