## Supplemental data for "Extracellular vesicles mediate the intercellular exchange of nanoparticles"

### Supplementary Information for
### Extracellular vesicles mediate the intercellular exchange of nanoparticles

Xian Wu[1], Tang Tang[1], Yushuang Wei[1], Katherine A. Cummins[2], David K. Wood[2], Hong-Bo Pang[1],*

[1] Department of Pharmaceutics, University of Minnesota, Minneapolis, MN, USA

[2] Department of Biomedical Engineering, University of Minnesota, Minneapolis, MN, USA

#### Supplementary Methods

##### Preparation of different nanoparticles.

70 nm silver nanoparticles (AgNPs) with PVP coating (AgNP-PVP) were prepared using a previous reported method[1]. 0.375 g of $AgNO_3$ (Sigma cat. no. 209139-25G >99%) and 1.5 g PVP (polyvinylpyrrolidone, Sigma cat. no. 856568-500G MW ~55000) were dissolved in 150 mL ethylene glycol (BDH cat. no. BDH1125-1LP >99.99%).The mixture was then heated to 160 °C and incubated for 1 h. After cooling, the silver NPs was precipitated in a large amount of acetone followed by centrifugation at 1000 RCF for 10 min. The solids were redispersed in water with bath sonication.

30 nm AgNPs with citrate coating were synthesized using a previous method[1]. 2.5 L water was heated up to 540 °C, and 450 mg $AgNO_3$ and 500 mg Sodium citrate tribasic dihydrate was dissolved in 50 mL water, respectively. After the water boiled, $AgNO_3$ and Sodium citrate tribasic dihydrate were added sequentially. After the mixture turned to dark greenish muddy stage, the reaction was kept for another 30 min. After the mixture cooled down to room temperature, it was centrifuge at 7000 RCF. AgNPs were collected and redispersed in water with bath sonication.

50 nm gold nanoparticles (AuNPs) were prepared using a previous reported method. Briefly, a 495.7 ml aqueous solution of 50 mg $HAuCl_4$ (Thermo Fisher Scientific Inc., cat. no. G545) was prepared and heated to boiling. A 4.3 ml of 34 mM trisodium citrate (Sigma cat. no. S4641) solution was then added. Heat removal after 5 min was followed by allowing the solution to cool to RT. After centrifuging at 5000 RCF for 10 min, the solids were washed with DI $H_2O$ and were redispersed at O.D. 40.

NeutrAvidin (NA, Thermo Scientific) was modified for silver nanoparticle coupling by appending 5 kDa NHS-PEG-OPSS (Jenkem), where OPSS is ortho-pyridyl disulfide. NA (25 mg) was dissolved in 5 mL 10% glycerol water solution for an hour, then brought to the final concentration of 1× PBS by mixing with 10× PBS. NHS-PEG- OPSS, 6 mg, was dissolved in 0.5 mL glycerol/water for 15 sec then added to the NA solution. After 4 to 5 h the NA-OPSS was dialyzed against 0.1× PBS with 2 mM NaN3 using a 20 kDa Slide-a-Lyzer, Pierce, 4 L buffer with one change after several hours followed by overnight dialysis. The product was filtered with a 0.22 μm syringe filter (Millipore), and then assayed for

OPSS using reduction by TCEP (tris(2-carboxyethyl)phosphine, 0.5M solution pH 7.0, Sigma).

Then silver/gold NeutrAvidin nanoparticles (Ag-NA and Au-NA) were prepared. Briefly, 0.5 mL of 2.8 mg/mL NA-OPSS in 0.1× PBS was added to 25 mL of 70 nm AgNP- PVP (O.D. 200) / 30 nm AgNP (O.D. 670) / 50 nm AuNP (O.D.40) in water and incubated overnight, then washed by centrifugation at 3300 RCF for 70 nm AgNP / 14000 RCF for 30 nm AgNP / 5000 RCF for 50 nm AuNP, decanting supernatant and sonicating into PBST. Lipoic PEG amine (MW 3400 g/mol, Nanocs Inc.) was dissolved in 70% ethanol and added to the Ag-NA (at a final concentration of 20 μM for 70 nm AgNP and 50 nm AuNP, 100 μM for 30 nm AgNP) and incubated for 3 days at room temperature. After washing and redispersing the particles in PBST, NA content was determined through biotin-4-fluorescein assay (Life Technologies, cat. no. B-10570) to be 102-103 NA per NP. NHS-CF647 (Biotium, cat. no. 92135) were added to portions of the Ag-NA/Au-NA at 20 μM final concentration, incubated overnight at 4 °C followed by extensive centrifuge washing with PBST until supernatant was not fluorescent. Biotin peptides (Biotin-TAT, Biotin-RPARPAR and Biotin-iRGD, LifeTein) were added at 40 μM for more than 1 h, then the solids were centrifuged, washed and redispersed with PBST.

Liposome and iRGD-Liposome were prepared as described previously. Briefly, DPPC, Cholesterol, and DSPE-PEG2000-Mal were weighed and dissolved with chloroform under a molar ratio of 16:4:10. The chloroform was removed by rotary evaporation and the film was hydrated with 10 mL HBS (10 mM, pH 7.4) solution containing 10% sucrose by using the rotavapor under 40 °C water bath condition but at normal atmosphere for around two hours. The primary liposomes were extruded with 1 μm, 0.2 μm and 0.1 μm membrane sequentially. For iRGD conjugation, proper molar of FAM-cysteine-iRGD was added into the solution to conjugate the maleimide on the surface of the liposomes. Unreacted FAM-Cysteine-iRGD was removed by dialysis with a 50kDa kit.

T-Dextran was prepared by conjugate TAT peptide with Dextran-tetramethylrhodamine (70,000 MW, lysine fixable, Fisher, D1818). Breifly, proper molar of NHS-PEG2000-Mal (Jenkem) was added into the solution of Dextran-tetramethylrhodamine to conjugated the maleimide on the surface of the Dextran. Then Cysteine-TAT (Lifetein) was then conjugated onto the Dextran through maleimide. Unreacted NHS-PEG2000-Mal and Cysteine-TAT were removed by dialysis with a 50kDa kit.

**Optimization of the intercellular exchange assay:**

To optimize the conditions for intercellular exchange assay, T-AgNPs was applied to PC-3 / PC3-GFP combinations in the assay. The transfer time (24 h, 48 h), cell number for recipient cells ($6\times10^4$, $9\times10^4$ and $1.2\times10^5$), donors to recipients ratio (2:1, 1:1, 1:2), concentration of gap collagen solution (2, 4 and 8 mg/mL), w / wo FBS in medium and the form of donor cells (monolayer or in collagen) was tested to get the optimal conditions.

We first tested the transfer time. $6\times10^4$ PC3-GFP cells were seeded in recipient layer as described in Methods. Then we added PC-3 cells into each well with the ratio of 1:1 and incubated for 24 h or 48 h. The apoptosis analysis was carried out with Dead Cell Apoptosis Kit (Thermo Fisher Scientific Inc., cat. no. V13241) followed by flow cytometry analysis. Data in Fig. S3A revealed that 48 h incubation led to a significant increase of cell apoptosis when compared to 24 h incubation. 24 h transfer time was chosen to all other experiments in this work.

Next, we optimized the recipient cell number and the ratio of donor cells to recipient cells in the intercellular exchange assay. $6\times10^4$ (60K in Fig. S3), $9\times10^4$ (90K in Fig. S3) and $1.2\times10^5$ (120K in Fig. S3) PC3-GFP cells were seeded in recipient layer as described in Methods, respectively. Then PC-3 cells was added into each well according to the ratio of 2:1, 1:1 and 1:2 (PC-3:PC3-GFP), respectively. After 24 h incubation, the intercellular exchange efficacy was tested and analyzed as described in Methods. As shown in Fig. S3B, the percentage of AgNP positive recipient cells increased as the ratio of donors to recipient increased regardless which number of recipient cells we were using. We also tried 6:1 (PC-3:PC3-GFP) ratio and no further increase of intercellular exchange efficacy was observed (Data not shown). Besides, 90K recipient cells group showed significant increase of AgNP positive recipient cells in the system than that of 60K group. Further increased the recipient cells to 120K didn't further increase this percentage. We then chose 90K recipient cells with 2:1 ratio of donor : recipient cells as the condition to do the rest of our study.

Whether the concentration of gap collagen solution between donor and recipient cells had any effect on the intercellular exchange was also investigated. The intercellular exchange assay was prepared as described in Methods. 2, 4, 8 mg/mL of collagen solution was used to prepare the gap between donor and recipient cells, respectively. The intercellular exchange efficacy was measured as described in Methods. As demonstrated in Fig. S3C, no significant difference of the intercellular exchange efficacy was observed among three groups. 2 mg/mL of collagen solution was used in our study if otherwise indicated.

Next, the effect of FBS in the medium on the intercellular exchange efficacy was tested. The assay was carried out in medium w / wo FBS, respectively. The intercellular exchange efficacy was measured as described in Methods. As demonstrated in Fig. S3D, no significant difference of the intercellular exchange efficacy was observed. Complete medium with FBS was used in all other intercellular exchange assays in this work.

Last, the effect of the format of donor cells on the intercellular exchange efficacy was investigated. The recipient layer and gap in the assay were prepared as described in Methods. Then we plated donor cells on top either as monolayer or encapsulated in a separate collagen layer and incubated for 24 h. Little difference in the intercellular exchange efficiency was observed between two groups (Fig.S3E). For simplicity, we used the monolayer format for donor cells in all other experiments.

**Confirmation of no direct cell-cell contact in the intercellular exchange assay:**

Intercellular exchange assays with different volumes of collagen gap (equivalent to 0,30 and 80 μL in 96-well plate) were prepared in 8 Chamber Polystyrene Vessel Tissue Culture Treated Glass Slide (Falcon, cat. no. 354108) as described in Methods. After 24 h incubation, cells were fixed with 500 μL 10% formalin overnight at room temperature. Formalin was removed and 800 μL O.C.T. compound (Tissue-Tek, SAKURA®, cat. no. 4583) was added into the chamber. Then the whole chamber was transferred into -80 °C overnight. The next day, the frozen bulk gel in the chamber was removed and frozen sections of bulk gel was prepared according to standard cryo-section protocol. Haematoxylin was applied to stain nuclei of cells in bulk gel using the standard method for H&E stain. Images were taken using EVOS M5000 microscope (Thermo Fisher Scientific). For immunofluorescence staining, PC3-GFP cells were used as donor and recipient cells. The frozen sections were prepared as described above. The chamber slides were then mounted in DAPI-containing mounting medium (Vector Laboratories, Burlingame, CA) with a coverslip and examined using EVOS M5000 microscope (Thermo Fisher Scientific).

**Evaluation of cell viability with GW4869 treatment:**

CellTiter-Glo® 3D Cell Viability Assay (Promega, catalog #: G9681) was applied to evaluate cell viability in 3D intercellular exchange assay. 3D intercellular exchange assay was prepared as described in Methods. After 24 h incubation, the cell viability was measured according to manufacturer's instruction of CellTiter-Glo® 3D Cell Viability Assay.

3-(4,5-dimethylthiazol-2-yl)-2,5-diphenyltetrazolium bromide (MTT, ) was applied to evaluate cell viability with GW4869 treatment. Cells was seeded at the density of $1.5\times10^4$ cells/well in a 96-well plate. After incubation for 24 h, the medium was changed to GW4869-containing medium at the dosage of 0, 10, 20 and 40 μM and cells were incubated for another 24 h. The medium was then discarded and 100 μL of 0.5 mg/mL MTT in FBS-free medium was added. After incubation at 37°C for 4 h, 150 μL DMSO was added into each well and mixed well by pipetting. The plate was incubated at room temperature for 30 min and then absorbance was measured at 570 nm using a SpectraMax M2 plate reader (Molecular Devices, Inc.).

**Evaluation of etching efficiency**

CPP-AgNPs was dispersed into PBS (O.D. 40). Equivalent volume of etchant concentration was added to CPP-AgNPs solution. A laser point was used to demonstrated the Tyndell effect[2,3] of the solution.

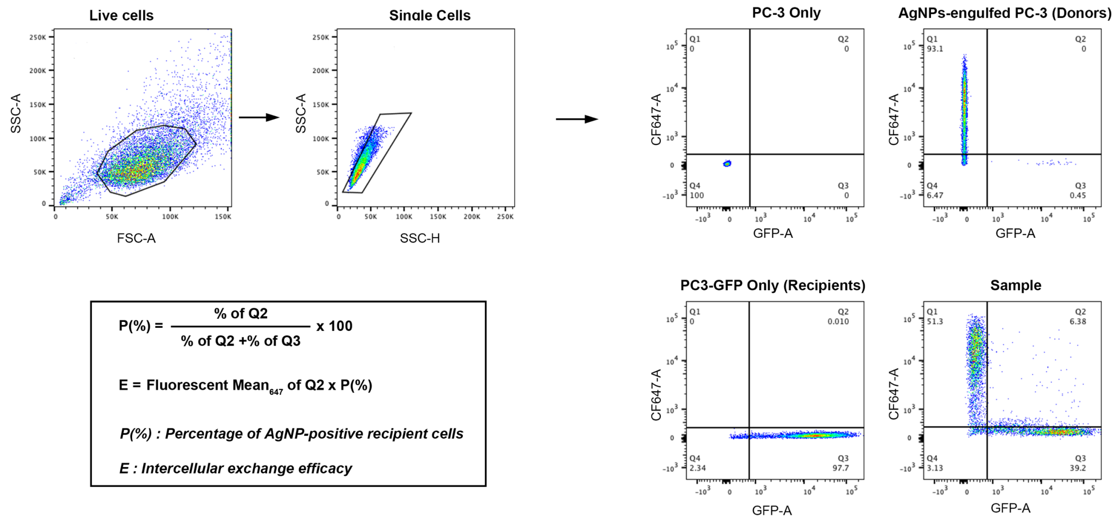


**Fig S1. The gating strategy for flow cytometry analysis and equations for calculating intercellular exchange efficacy in this work.**

Example of gating strategy for flow cytometry analysis. After the intercellular exchange assay was completed, cells were isolated, fixed and analyzed using a BD FACS Calibur flow cytometry as described in Methods. FlowJo was used to analyze the data. Single cells were picked out followed by quad gating for AgNP-positive recipient cells (Q2) based on different single-stain controls (PC-3 Only, AgNP-engulfed PC-3 and PC3-GFP Only). The intercellular exchange efficacy was calculated using equations showed in the box.

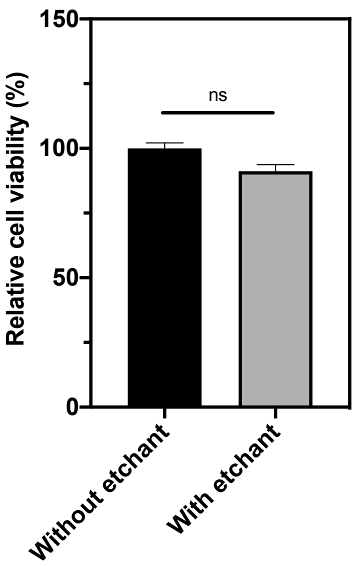


**Fig S2. Cell viability under constant etching process in 3D intercellular exchange assay.**

Cell viability of PC-3 (donors) and PC3-GFP (recipients) in 3D intercellular exchange assay after 24 h incubation with presence or absence of constant etching (x axis) was tested as described in Supplementary Methods, respectively. The result was normalized to that of without etching (y axis). Error bars indicate S.E.M., n=3.

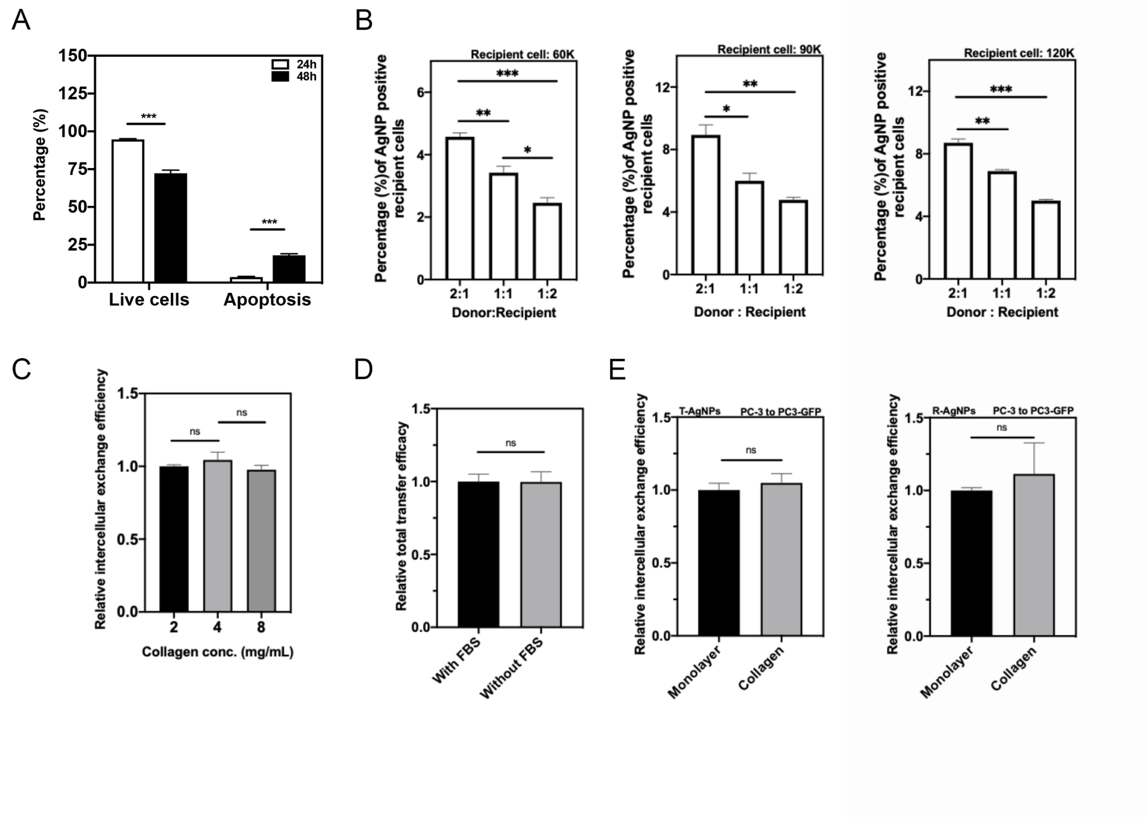


**Fig. S3 The optimization of intercellular exchange assay.**

(A) Quantitative analysis of apoptosis of cells in the intercellular exchange system after 24 h and 48 h. (B) Percentage of AgNP positive recipient cells (y axis) with different recipient cell numbers and ratio of donor : recipient (x axis) after 24 h was quantified as described in Supplementary Methods. (C) Intercellular exchange efficacy of T-AgNPs with indicated collagen concentration for the gap (x axis) after 24 h was quantified as described in Supplementary Methods, result was normalized to that of with 2 mg/mL (y axis). (D) Intercellular exchange assay was carried out in medium w / wo FBS as described in Supplementary Methods, respectively. Intercellular exchange efficacy of T-AgNPs w / wo FBS in the medium (x axis) after 24 h was quantified and normalized to that of with FBS in the medium (y axis). (E) Intercellular exchange efficacy with different format of donor cells for T-AgNPs and R-AgNPs. Intercellular exchange efficacy of T-AgNPs and R-AgNPs from PC3 to PC3-GFP cells was quantified as described in Methods and normalized to that of monolayer condition (y-axis). Error bars indicate S.E.M. (standard error of the mean), n=3. *P<0.05, **P<0.01, ***P<0.001 and ns, no significance (Student's t-test).

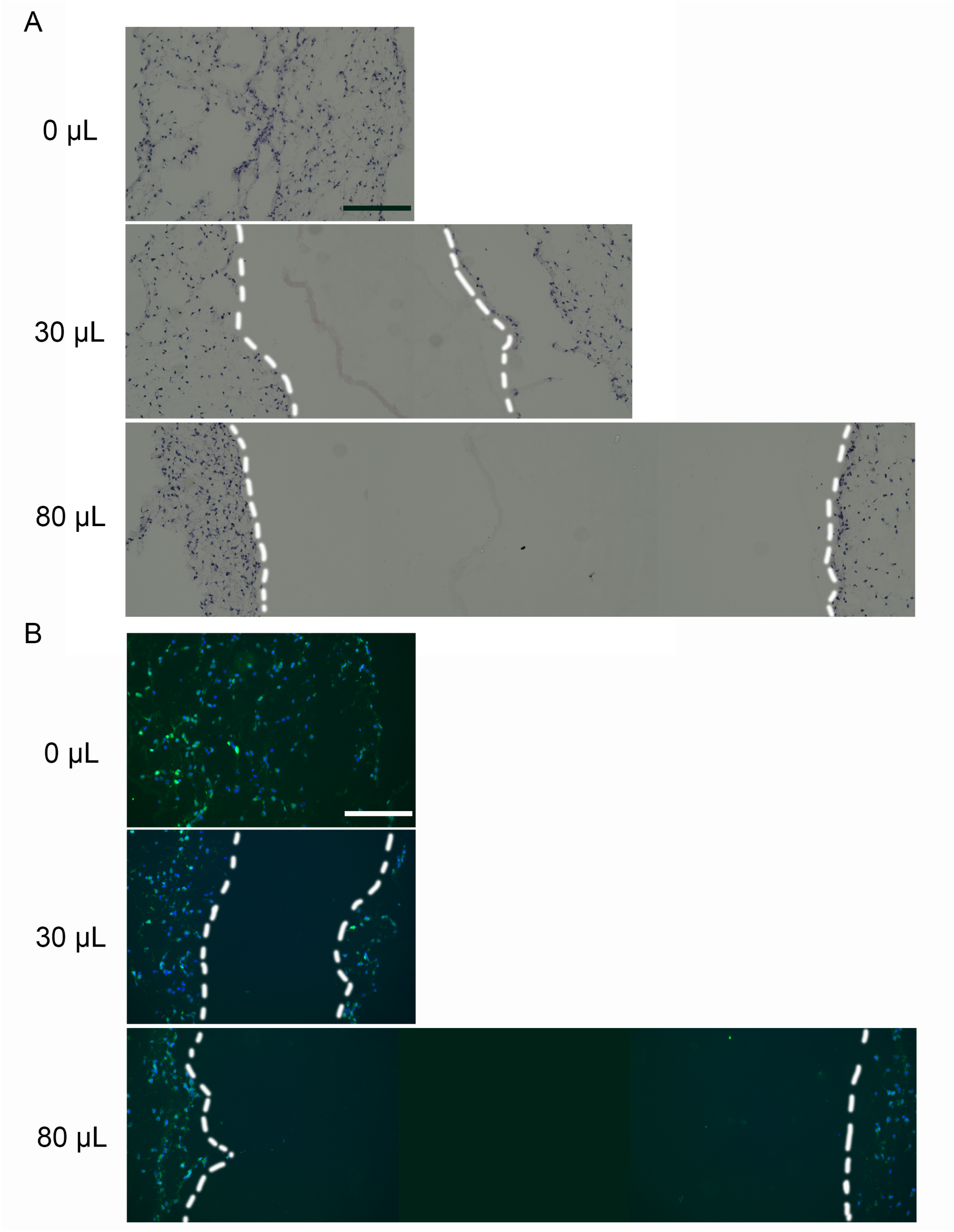


**Fig. S4 No direct cell-cell contact in the intercellular exchange assay with collagen gap.** Immune-histochemical (A) and immunofluorescence (B) staining of PC3-GFP cells on both sides of collagen gaps with indicated volume of collagen solution in the intercellular exchange assay. All slides were prepared and imaged as described in Supplementary Methods. White dash line indicated the boundary of cells that seeded on both sides of collagen gaps. At least three independent experiments were performed and representative images are shown here. Scale bar, 300 μm.

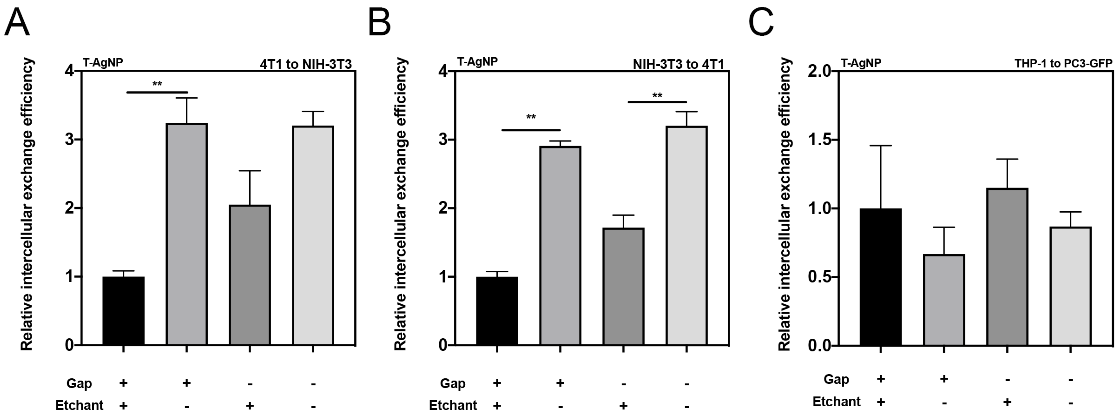


**Fig. S5 Intercellular exchange between other pairs of donor and recipient cells.**

Intercellular exchange efficacy of T-AgNPs from 4T1 to NIH-3T3 (A), NIH-3T3 to 4T1 (B) and THP-1 to PC3-GFP (C) under the indicated conditions (x axis) was measured as described in Methods and normalized to that of group under constant etching with collagen gap (y axis). Error bars indicate S.E.M., n=3. **P<0.01 (Student's t-test).

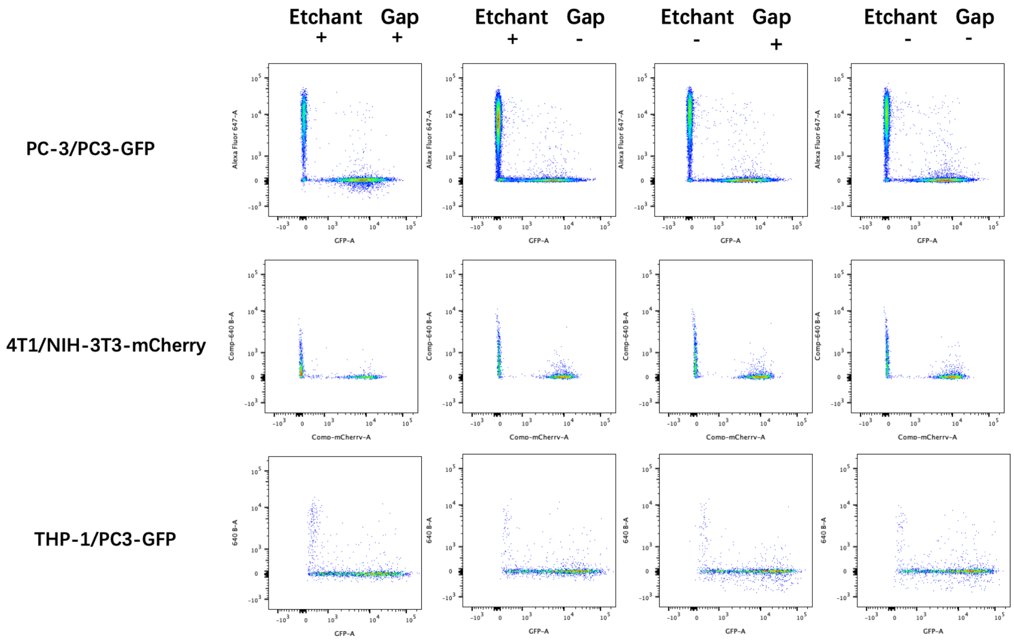


**Fig. S6 Representative flow cytometry data of intercellular exchange assay.**

3D intercellular exchange assays of T-AgNPs with indicated donor / recipient combinations was carried out and tested by flow cytometry as described in Methods. Data was analyzed as showed in Fig. S2.

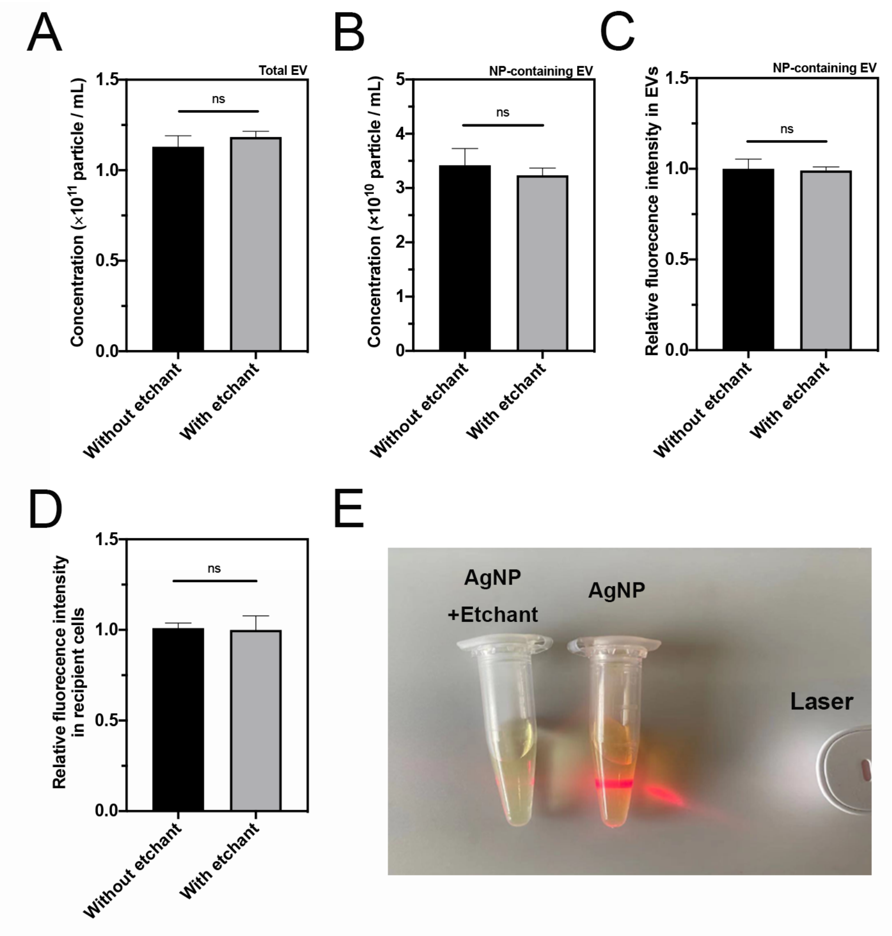


**Fig. S7 Etching process didn't impair the capacity of EV secretion from donor cells nor EV uptake byrecipient cells.**

(A-C) Etching process showed little effect on EV secretion from donor cells. Total EVs (A) and NP-containing EVs (B) from PC-3 cells after 24 h incubation w / wo etching (x axis) was isolated and the particle number (y axis) was measured by NTA as described in Methods. (C) NP-containing EVs from PC-3 cell after 24 h incubation w / wo etching (x axis) was isolated, the relative fluorescence intensity inside EVs was measured as described in Methods. The result was normalized to that of without etching (y axis). (D) Etching process showed little effect on EV uptake by recipient cells. NP-free EVs from PC-3 cell after 24 h incubation w/wo etching (x axis) was isolated and labeled with Dil as described in Methods. Labeled EVs were incubated with PC3-GFP cells for 30 min and the relative fluorescence intensity of Dil in PC3-GFP cells was measured by flow cytometry and normalized to that of without etching (y axis). (E) The concentration of etchant used in this study is effective for eliminating free AgNPs in the assay.

Tyndall effect disappeared after equivalent volume of etchant was added to AgNP solution indicated by the loss of scattering of the laser. Error bars indicate S.E.M., n=3. ns, no significance (Student's t-test).

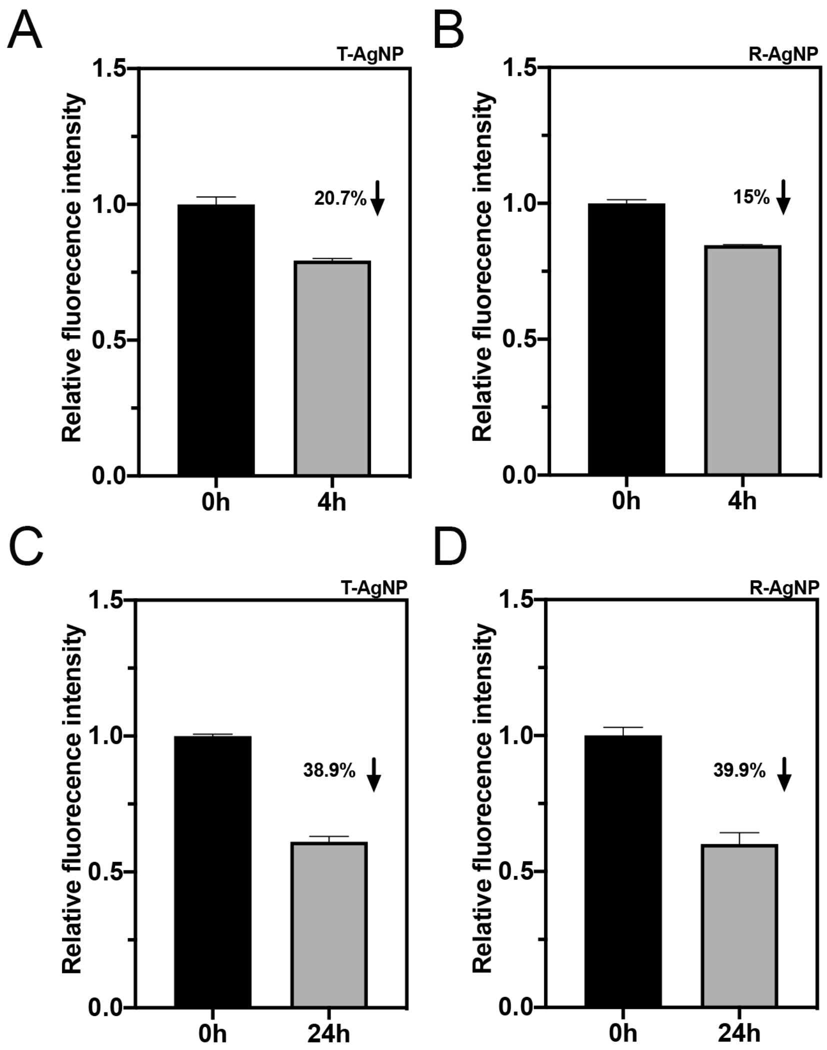


**Fig. S8 Endocytic and exocytic profile of R-AgNP and T-AgNP by PC-3 cells.**

(A and B) Uptake of CPP-AgNPs by donor cells. PC-3 cells as donor cells were incubated with T-AgNP / R-AgNP for 4 h and the fluorescence intensity of T-AgNP (A) and R-AgNP (B) in the culture medium (4 h) was measured and compared with that of the initial input into the well (0 h). (C and D) exocytosis of CPP-AgNPs by donor cells. PC-3 cells as donor cells were incubated with T-AgNP / R-AgNP for 4 h and washed with etchant as described in Methods. The intracellular fluorescence intensity was measured by flow cytometry (0h). After 24 h incubation in fresh medium, the intracellular fluorescence intensity was measured again by flow cytometry (24 h). Error bars indicate S.E.M., n=3.

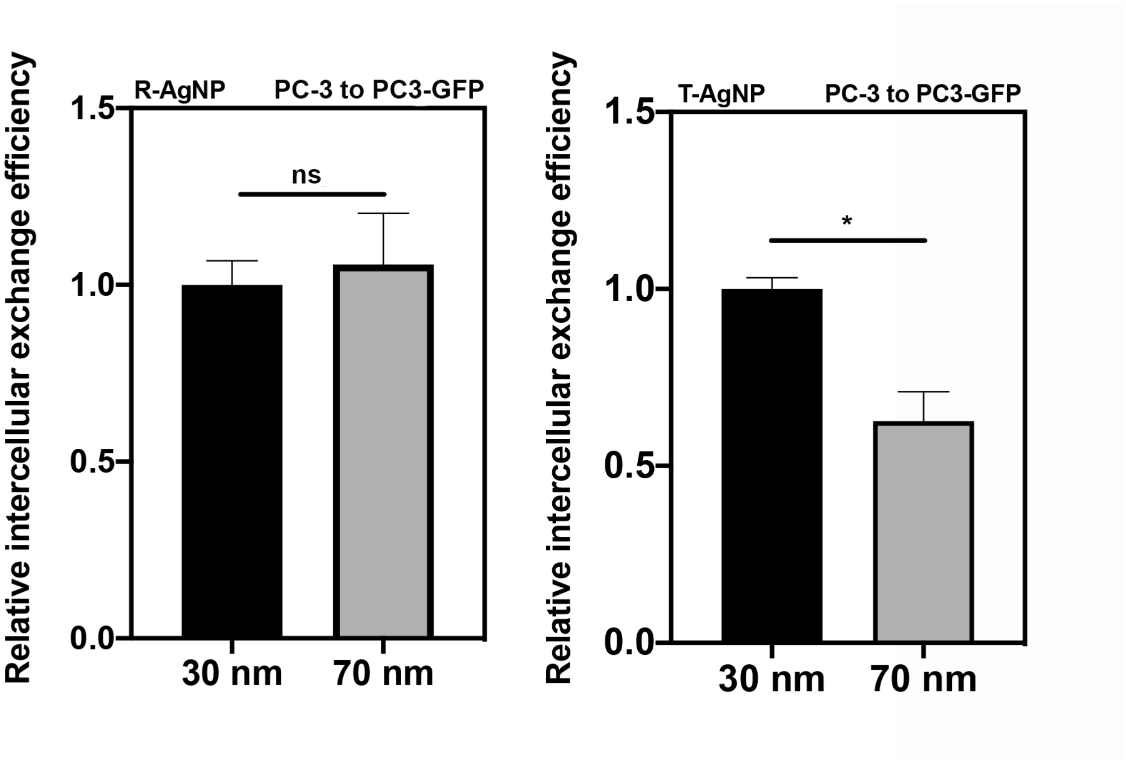


**Fig. S9 Intercellular exchange efficacy with different size of CPP-AgNPs in PC-3/PC3-GFP cells.**

The intercellular exchange efficacy of 30 nm and 70 nm T-AgNPs / R-AgNPs from PC-3 to PC3-GFP cells was quantified as described in Methods and the result was normalized to that of 30 nm CPP-AgNPs (y axis). Error bars indicate S.E.M., n=3. ns, no significance (Student's t-test).

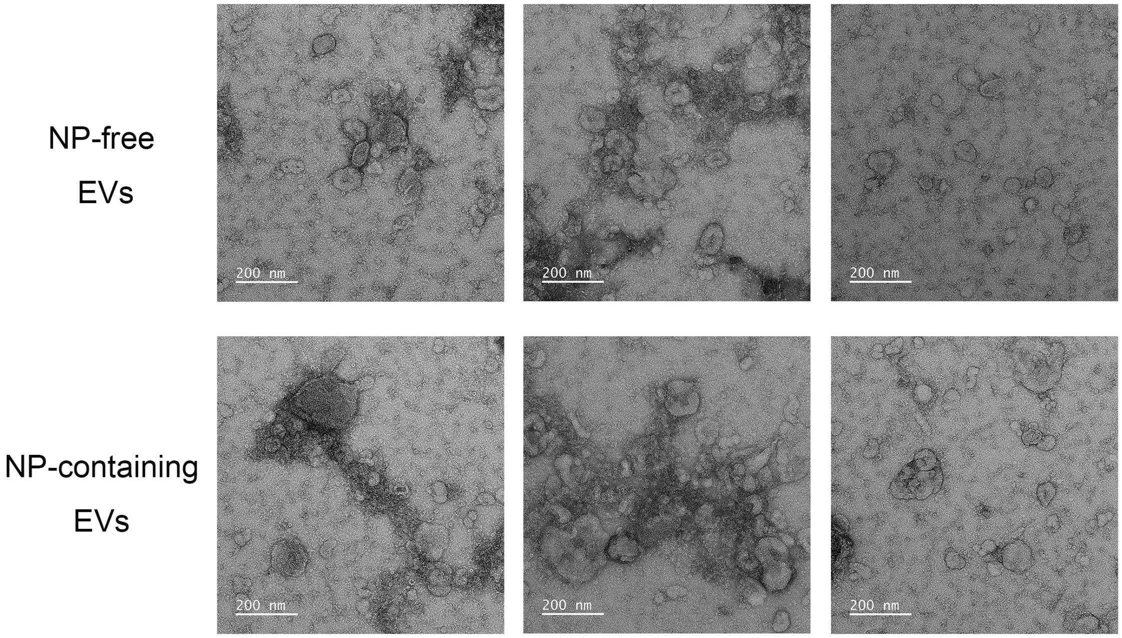


**Fig. S10 Representative TEM images of NP-free EVs and NP-containing.**

NP-free EVs and NP-carrying EVs were isolated from PC-3 cells pre-incubated with T-AgNPs by density gradient ultracentrifugation as described in Methods. Representative TEM images of NP-free EVs and NP-containing EVs. Scale bar, 200nm.

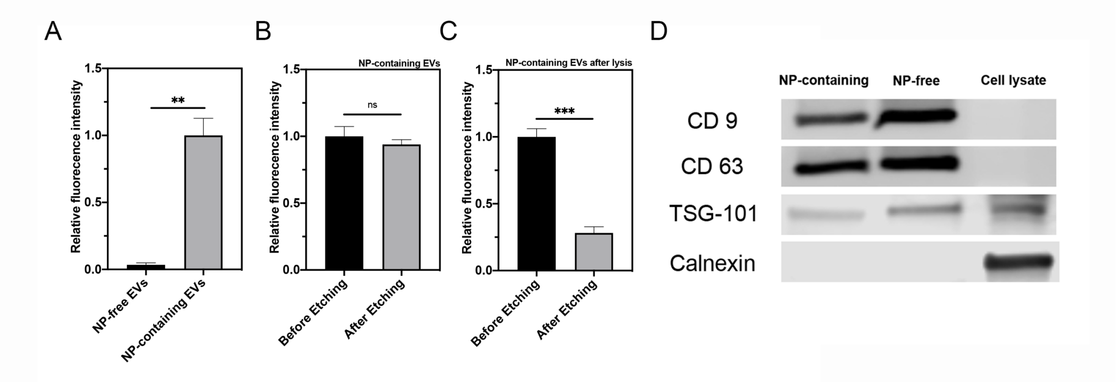


**Fig. S11 Validation of R-AgNPs exocytosed inside HUVEC-derived EVs.**

R-AgNPs were exocytosed by EVs from HUVEC cells. NP-free EVs and NP-containing EVs were isolated from HUVEC cells pre-incubated with R-AgNPs by density gradient ultracentrifugation and characterized as described in Methods. (A) Fluorescence intensity of R-AgNPs in NP-free EVs and NP-containing EVs was detected and normalized to that of in NP-containing EVs (y axis). (B and C) Before (B) and After (C) treating NP-containing EVs with TX-100 detergent, the fluorescence intensity of R-AgNPs within before / after etching was detected and the result was normalized to that of before etching (y axis). (D) Western blot analysis of NP-free and NP-containing EVs. The present of canonical exosome markers, including CD9, CD63 and TSG-101 and the absence of Calnexin were detected from NP-free EVs and NP-containing EVs. Error bars indicate S.E.M., n=3. **P<0.01, **** P<0.0001 and ns, no significance (Student's t-test).

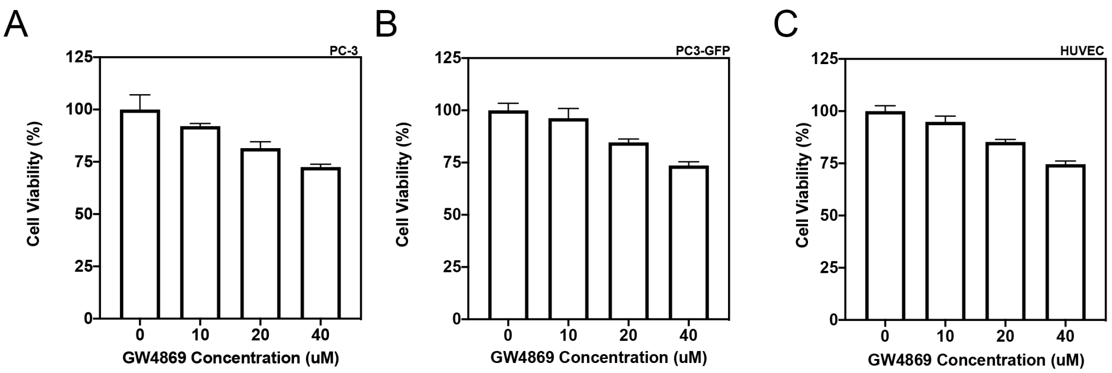


**Fig. S12 Cell viability analysis with GW4869 treatment.**

Cell viability of PC-3 (A), PC3-GFP (B) and HUVEC (C) after 24 h incubation with GW4869 at indicated concentrations (x axis) was tested by standard MTT assay as described in Supplementary Methods, respectively. The result was normalized to that of without GW4869 treatment (y axis). Error bars indicate S.E.M., n=5.

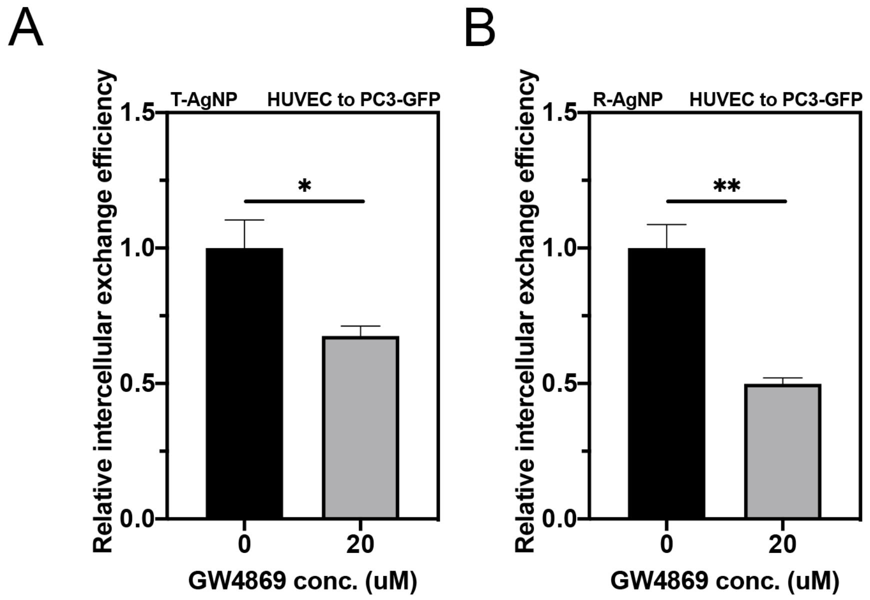


**Fig. S13 GW4869 decreased intercellular exchange of CPP-AgNPs from HUVEC to PC3-GFP cells.**

Intercellular exchange efficacy of T-AgNPs (A) and R-AgNPs (B) from HUVEC to PC3-GFP cells with indicated GW4869 concentrations (x axis) was quantified as described in Methods and normalized to that of without GW4869 treatment (y axis). Error bars indicate S.E.M., n=3. *P<0.05 and **P<0.01 (Student's t-test).

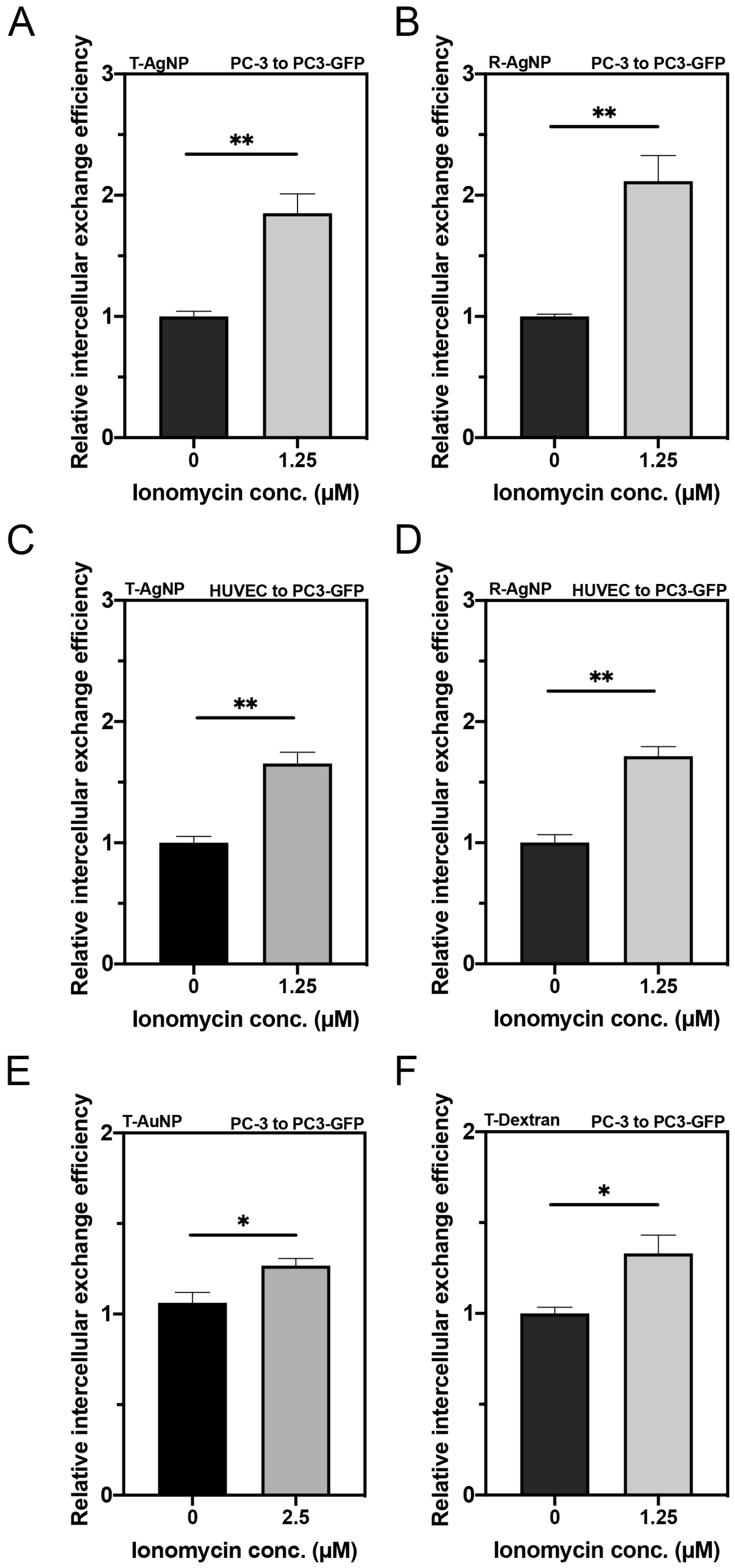

A
T-AgNP
PC-3 to PC3-GFP
Relative intercellular exchange efficiency
**
Ionomycin conc. (μM)
B
R-AgNP
PC-3 to PC3-GFP
Relative intercellular exchange efficiency
**
Ionomycin conc. (μM)
C
T-AgNP
HUVEC to PC3-GFP
Relative intercellular exchange efficiency
**
Ionomycin conc. (μM)
D
R-AgNP
HUVEC to PC3-GFP
Relative intercellular exchange efficiency
**
Ionomycin conc. (μM)
E
T-AuNP
PC-3 to PC3-GFP
Relative intercellular exchange efficiency
*
Ionomycin conc. (μM)
F
T-Dextran
PC-3 to PC3-GFP
Relative intercellular exchange efficiency
*
Ionomycin conc. (μM)

**Fig. S14 Ionomycin treatment increased the intercellular exchange of CPP-NPs.**

(A-D) Ionomycin increased intercellular exchange of CPP-AgNPs in different donor/recipient pairs. Intercellular exchange efficacy of T-AgNPs (A) and R-AgNPs (B) from PC-3 to PC3-GFP cells with indicated ionomycin concentrations (x axis) was quantified as described in Methods and normalized to that of without ionomycin treatment (y axis). Intercellular exchange efficacy of T-AgNPs (C) and R-AgNPs (D) from HUVEC cells to PC3-GFP cells with indicated ionomycin concentrations (x axis) was quantified as described in Methods and normalized to that of without ionomycin treatment (y axis). (E and F) Ionomycin increased intercellular exchange of other types of CPP-NPs. Increased intercellular exchange efficacy of T-AuNP (E) and T-Dextran (F) from PC-3 to PC3-GFP cells with indicated ionomycin concentrations (x axis) was quantified as described in Methods and normalized to that of without ionomycin treatment (y axis).Error bars indicate S.E.M., n=3. *$P<0.05$ and **$P<0.01$ (Student's t-test).

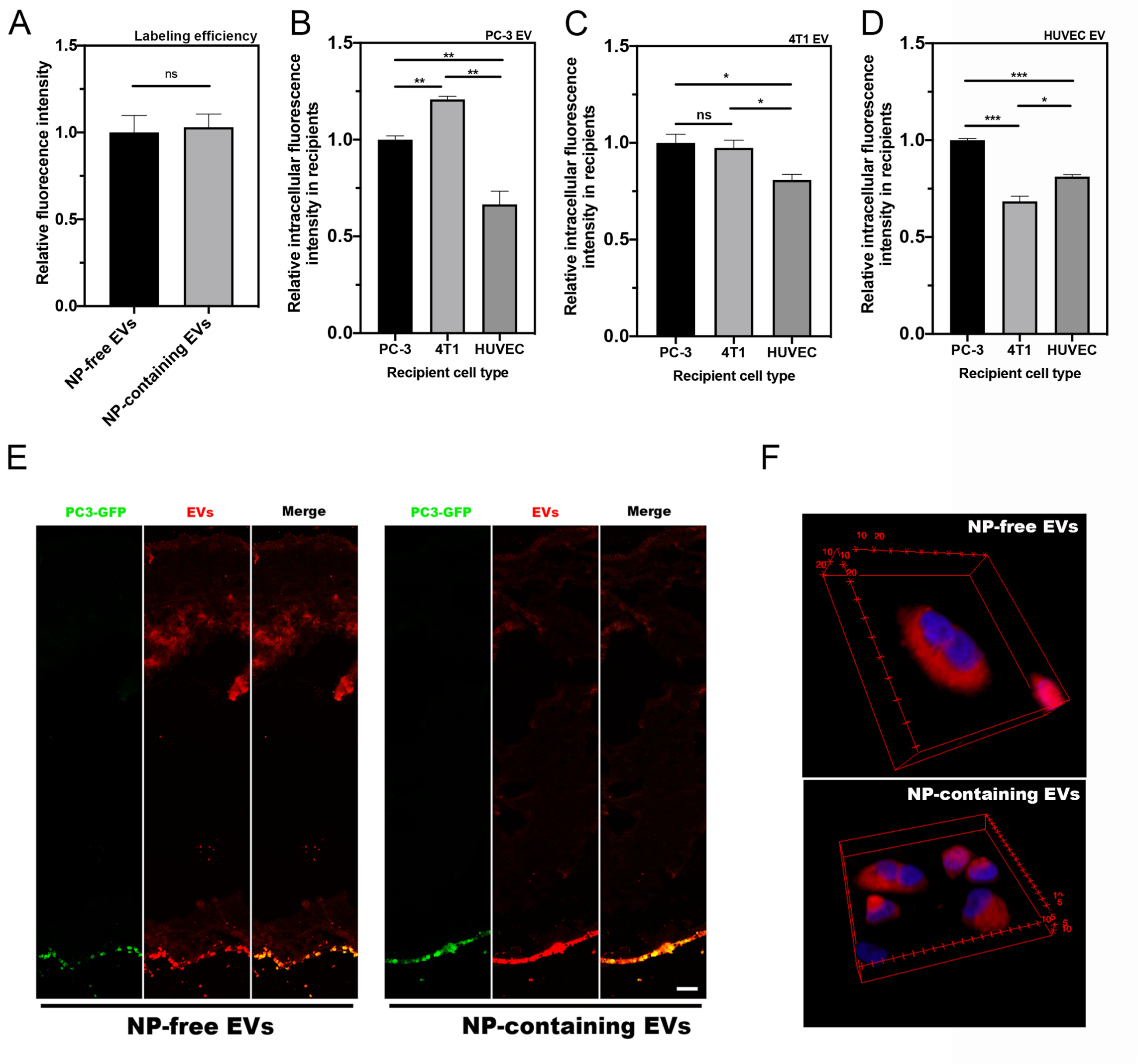


**Fig. S15 Re-entry of NP-containing EVs into recipient cells.**

(A) Labeling efficiency of Dil is similar for NP-free and NP-containing EVs. NP-free and NP-containing EVs were isolated from T-AgNPs engulfed PC-3 cells. Same particle number of two EV subsets (determined by NTA) was labeled with Dil as described in Methods. The fluorescence intensity of two subsets (x axis) was measured and normalized to that of NP-free EVs (y axis). (B-D) Cell entry efficiency of EVs depends on both parent and recipient cell types. EVs were isolated from the indicated cell types. After Dil labeling and normalization by protein content, EVs were incubated with the indicated recipient cells for 2 h before washing and imaging (x-axis). The intracellular EVs were quantified as described in Methods and normalized to that of PC-3 cells (y-axis). (E) Penetration of NP-free and NP-containing EVs through collagen matrix. NP-free and NP-containing EVs were isolated from T-AgNPs engulfed PC-3 cells, labeled with Dil (red) and added to the top of a collagen matrix (2 mg/mL collagen concentration) with a layer of PC3-GFP cells (green) underneath to mark the bottom. Both NP-free EVs (left panel) and NP-containing (right panel) showed co-localization with PC3-GFP cells after 30 min incubation at 37 °C. At least three independent experiments were performed and representative images are shown here. Scale bar, 100 μm. (F) Representative images of confocal microscopy with 3D reconstruction of cells after incubated with DiL labeled EVs. NP-free and NP-containing EVs were isolated from T-AgNPs engulfed PC-3 cells, labeled with Dil (red) and incubated with PC-3 cells for 30 min, respectively. 52 slides of confocal images were taken for each view and 3D reconstruction was carried out with ImageJ. Error bars indicate S.E.M., n=3. *P<0.05, **P<0.01, *** P<0.001 and ns, no significance (Student's t-test).

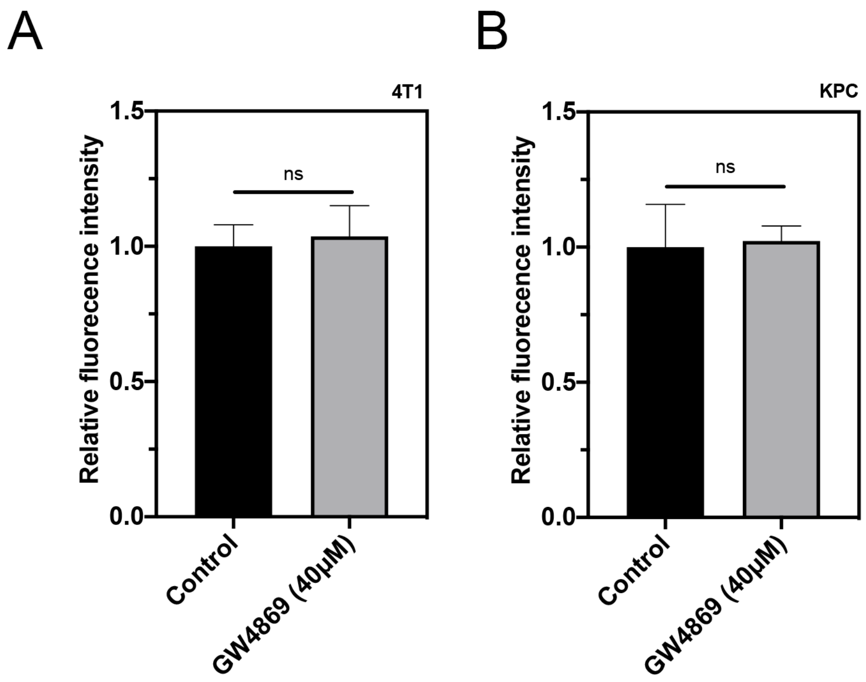


**Fig. S16 GW4869 induced little toxicity in tumor tissue.**

4T1 and KPC tumor model were established and GW4869 was administrated as described in Methods. Tumor was excised and sectioned for Tunel staining using In Situ Cell Death Detection Kit, Fluorescein (Sigma, cat# 11684795910). Quantitative analysis of fluorescence intensity of Tunel signal in 4T1 tumor (A) and KPC tumor (B) by ImageJ and normalized to that of control group (y axis). The experiment was performed in 3 mice per group. 3 images from each tumor tissue were used to analyze Tunel signal intensity in each group. Error bars indicate S.E.M.. ns, no significance (Student's t-test).

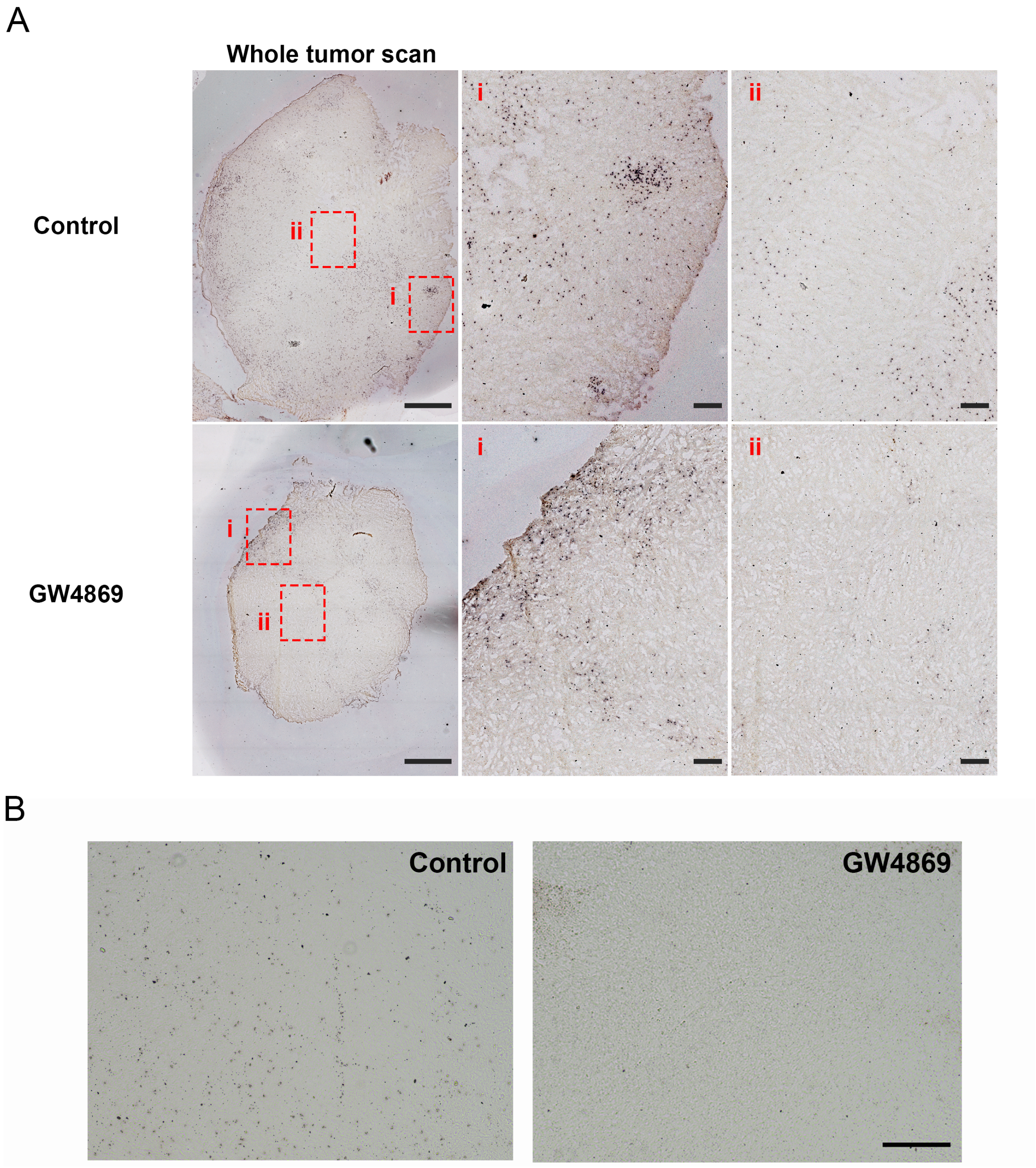


**Fig. S17 Representative images of Ag signal in 4T1 orthotopic tumor model and orthotopic pancreatic ductal adenocarcinoma tumor model after GW4869 treatment.**

GW4869 decreased the accumulation and penetration of iRGD-AgNPs in orthotopic 4T1 and pancreatic ductal adenocarcinoma tumor model *in vivo*. 4T1 tumor bearing mice received 20 μL of GW4869 (40 μM) via intratumoral injection each day for 5 days. Orthotopic pancreatic ductal adenocarcinoma tumor bearing mice received GW4869 at a dosage of 2.5 μg/kg body weight via intraperitoneal injection every other day for 5 injections. 24 h after the last injection, 50 µL of iRGD-AgNPs (O.D 40) was intravenously injected and circulated for 4 h. Tumor was excised and sectioned for AgNP and blood vessel detection as described in Methods. (A) Representative whole tissue scan images and zoomed areas (red squares) of 4T1 tumor. Scale bar, 1000 µm; insets, 200 µm. (B) Representative pictures of orthotopic pancreatic ductal adenocarcinoma tumor tissue with silver enhancement staining. Scale bar, 200 µm. The *in vivo* experiment was performed in 3 mice per group.

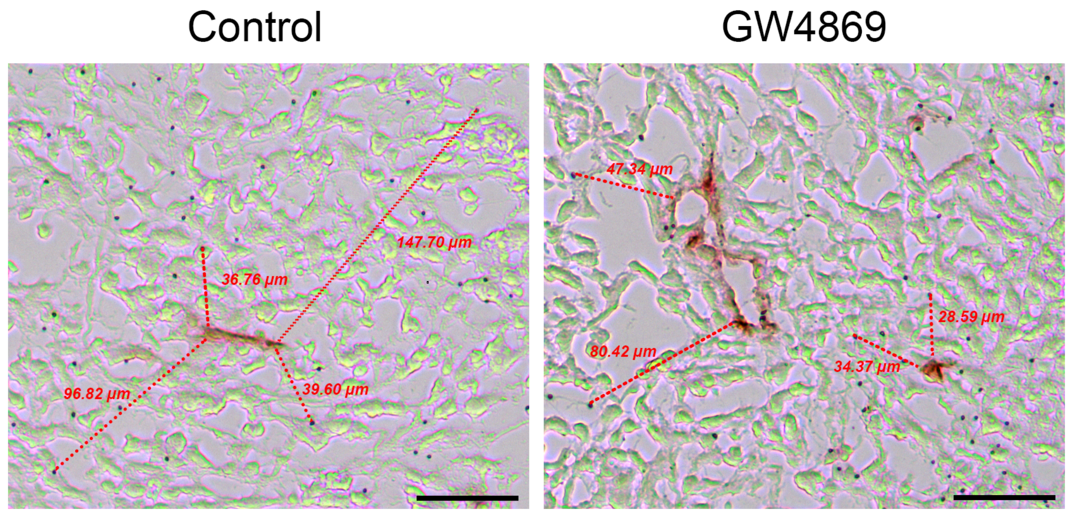


**Fig. S18 Analysis of penetration distance of iRGD-AgNPs from the nearest blood vessel in poorly vascularized region in 4T1 tumor model *in vivo*.**

Examples of analysis of penetration distance of iRGD-AgNPs from the nearest blood vessel in 4T1 tumor model. Scale bar, 50 µm. 4T1 tumor bearing mice received 20 µL of GW4869 (40 µM) via intratumoral injection each day for 5 days. 24 h after the last injection, 50 µL of iRGD-AgNPs (O.D 40) was intravenously injected and circulated for 4 h. Tumor was excised and sectioned for AgNP and blood vessel detection as described in Methods. The penetration distance was analyzed by ImageJ.
